# Mechanical stress compromises multicomponent efflux complexes in bacteria

**DOI:** 10.1101/571398

**Authors:** Lauren A. Genova, Melanie F. Roberts, Yu-Chern Wong, Christine E. Harper, Ace George Santiago, Bing Fu, Abhishek Srivastava, Won Jung, Lucy M. Wang, Łukasz Krzemiński, Xianwen Mao, Xuanhao Sun, Chung-Yuen Hui, Peng Chen, Christopher J. Hernandez

## Abstract

Physical forces have long been recognized for their effects on the growth, morphology, locomotion, and survival of eukaryotic organisms^1^. Recently, mechanical forces have been shown to regulate processes in bacteria, including cell division^2^, motility^3^, virulence^4^, biofilm initiation^5,6^, and cell shape^7,8^, although it remains unclear how mechanical forces in the cell envelope lead to changes in molecular processes. In Gram-negative bacteria, multicomponent protein complexes that form rigid links across the cell envelope directly experience physical forces and mechanical stresses applied to the cell. Here we manipulate tensile and shear mechanical stress in the bacterial cell envelope and use single-molecule tracking to show that shear (but not tensile) stress within the cell envelope promotes disassembly of the tripartite efflux complex CusCBA, a system used by *E. coli* to resist copper and silver toxicity, thereby making bacteria more susceptible to metal toxicity. These findings provide the first demonstration that mechanical forces, such as those generated during colony overcrowding or bacterial motility through submicron pores, can inhibit the contact and function of multicomponent complexes in bacteria. As multicomponent, trans-envelope efflux complexes in bacteria are involved in many processes including antibiotic resistance^9^, cell division^10^, and translocation of outer membrane components^11^, our findings suggest that the mechanical environment may regulate multiple processes required for bacterial growth and survival.

---

Physical forces play a fundamental role in the form and function of organs and organisms^1^. In eukaryotic systems, mechanical forces regulate morphogenesis and embryogenesis^12^, tissue healing and regeneration^13^, and the development of disease^14^. Much less is known about the importance of mechanical forces to prokaryotic organisms. Bacteria have long been known to respond to mechanical stresses associated with changes in osmotic pressure and hydrostatic pressure, but recent findings indicate that mechanical stresses caused during locomotion^3^, surface adhesion^4,5^, and cell division^2^ also regulate bacterial physiology.

Mechanical forces associated with adhesion to surfaces or locomotion can induce tensile (lengthening), compressive (shortening), and shear (shape changing) stresses to the inner, outer membranes and periplasmic region of the bacterial cell envelope, a combination of stresses that is much more complicated than those caused by changes in osmolarity. To query the individual contributions of each form of mechanical stress, we used a microfluidic device with submicron features to apply mechanical loads to individual bacteria. The device is analogous to micropipette aspiration commonly used to study mammalian cell biomechanics^15,16^ but instead of pulling the cell into a tapered channel, the device forces cells into tapered channels using fluid pressure^17^. Each device contains sets of tapered channels to apply twelve distinct magnitudes of pressure difference (Δ*P*) across the trapped bacteria within a single experiment (Fig. 1a, 1b; and Extended Data Fig. 1). The pressure difference is controlled by modifying fluid pressure at the inlet (also affecting the average pressure, *P*_ave_, which is indicative of hydrostatic pressure experienced by the cell) and determined locally with hydraulic circuit models (Supplementary Note 1.2). We refer to this loading modality as “extrusion loading.” Bacteria submitted to stepwise increases in Δ*P* exhibited increases in cell length and decreases in cell width, resulting in a net reduction in cell volume (Fig. 1c and Extended Data Fig. 2). Analytical and finite element models indicate that extrusion loading causes increases in axial tensile stress, reductions in hoop (transverse) tensile stress, and increases in shear stress, related to the magnitude of Δ*P* (Fig. 1d and Extended Data Fig. 3 and 4). Furthermore, analytical examination shows that reductions in cell volume during extrusion loading result in an increase in cell internal pressure, which we attribute to increases in osmolarity associated with loss of water from the cytoplasm when cell volume declines.

**Figure 1.**
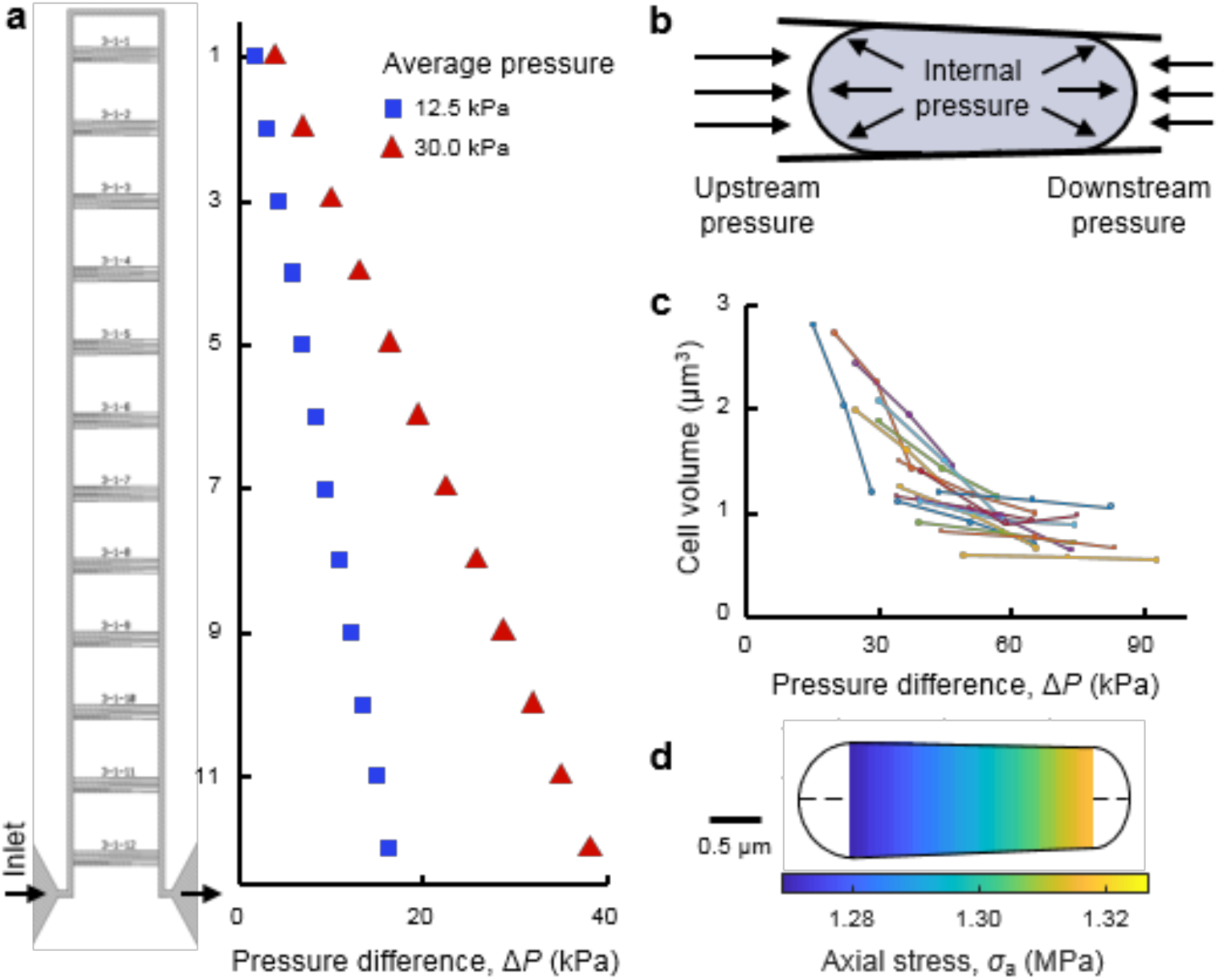
Mechanical loading of bacteria via a microfluidic device. **a**, A functional unit of the microfluidic device has twelve sets of five tapered channels. Fluid flow enters the functional unit at the bottom left, travels around the bypass channel, and exits out the bottom right. The difference between upstream and downstream pressure is larger for tapered channels closer to the inlet and outlet. Increasing the applied pressure increases the pressure difference Δ*P* at each set of tapered channels (and also increases the average pressure, *P*_ave_). **b**, Trapped bacteria experience greater upstream pressure than downstream pressure. The pressure difference Δ*P* is defined as the difference between the upstream and downstream pressures. Internal pressure due to turgor is also present. **c**, Increases in Δ*P* via stepwise increases of externally applied pressure results in reduced cell volume of trapped cells. Lines connect measurements of same cells. **d**, Analytical modeling of a trapped cell indicates a linear increase in axial stress along the length of the cell.

In eukaryotic organisms, sensitivity to mechanical forces is derived primarily from the transmission of forces from the extracellular environment into the cell through transmembrane protein complexes^18^. Many trans-envelope protein complexes exist in prokaryotes. In Gram-negative bacteria, protein efflux complexes from the resistant-nodulation-division (RND) family span the cell envelope and provide clinically relevant multidrug resistance by exuding toxic molecules from the cell^9^. To understand the effects of mechanical stress on bacterial efflux, we examined CusCBA, a tripartite Cu^+^ and Ag^+^ efflux complex of the RND family in *E. coli*. CusA is a trimeric proton-motive-force–driven pump located in the inner-membrane; CusB is a soluble periplasmic adaptor protein; CusC is a trimeric outer-membrane pore protein. These three proteins assemble into the complete CusC_3_B_6_A_3_ complex to enable efflux of Cu^+^/Ag^+^ from the cell^19-21^. In a cellular environment, CusCBA exists in a dynamic equilibrium between an assembled and disassembled state; upon increasing environmental copper concentration, the CusB protein binds copper and drives the complex toward the assembled state to enable copper efflux (Fig. 2c, inset)^22^.

**Figure 2.**
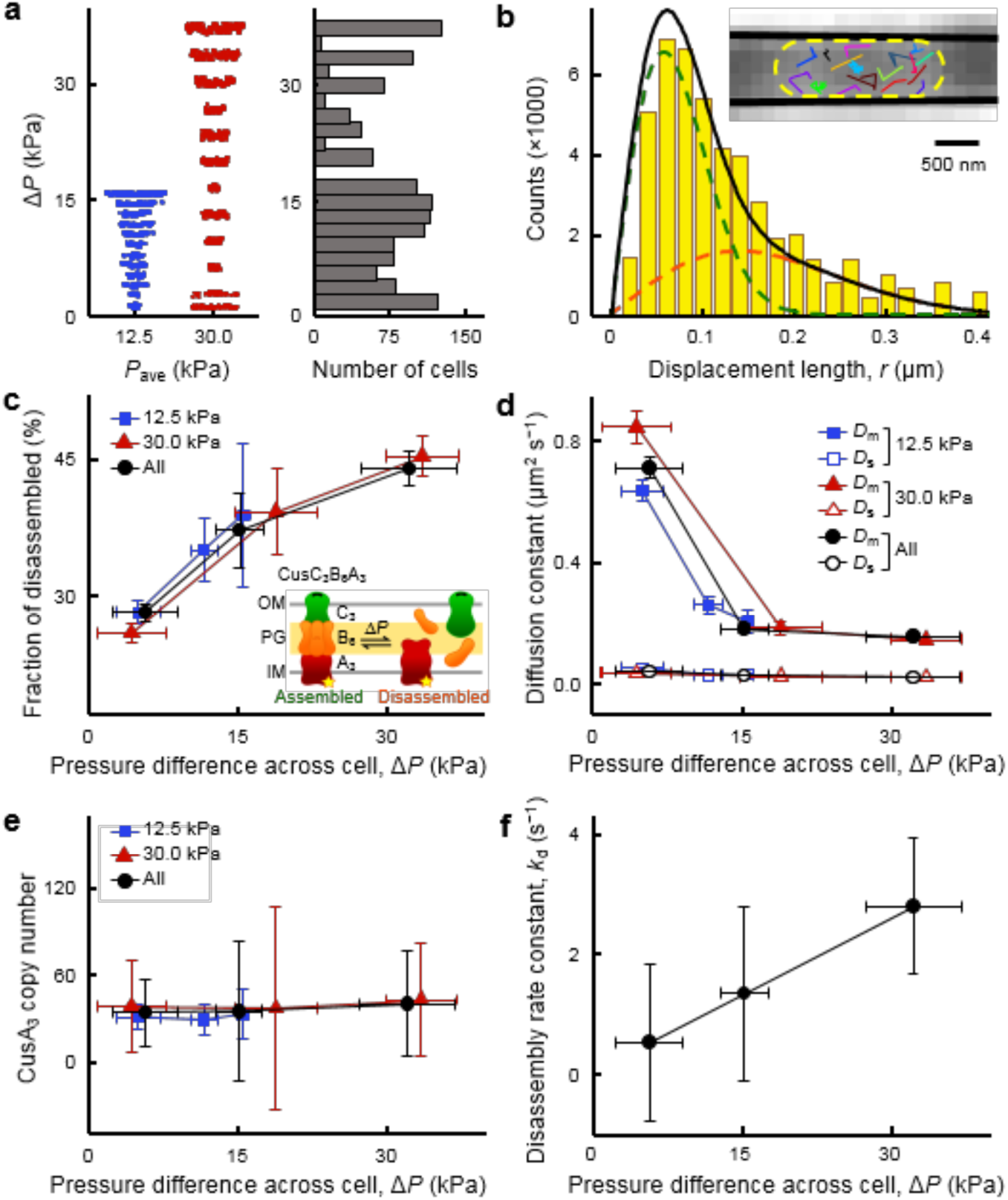
Single-molecule tracking uncovers mechanical stress–induced CusCBA disassembly. **a**, Cells were examined at two applied external loading conditions resulting in average pressure values of 12.5 kPa (*n* = 592 cells) and 30 kPa (*n* = 732 cells), giving a range of Δ*P* across individual cells. **b**, Distribution of displacement length *r* per time lapse for single CusA^mE^ proteins at *P*_ave_ = 30 kPa and Δ*P* = 32 ± 5 kPa, in which the cell confinement effect is deconvoluted (Supplementary Note 1.12). The distribution here resolves minimally two Brownian diffusion states (Eq. S23): a mobile disassembled state (orange dashed line) and an almost stationary assembled state (green dashed line), with diffusion constants of *D*_m_ = 0.16 ± 0.01 μm^2^ s^−1^ and *D*_s_ = 0.027 ± 0.001 μm^2^ s^−1^ and fractional populations of *A*_m_ = 44 ± 2% and *A*_s_ = 56 ± 2%, respectively. Solid black line: overall fit. Inset: Overlay of many position trajectories of single CusA^mE^ proteins in a living *E. coli* cell trapped in a tapered channel. Each colored line is from one CusA^mE^. Yellow dashed line: cell boundary; solid black lines: inner walls of the tapered channel. **c**, Fractional populations of the mobile disassembled state of CusA^mE^ increases with increasing Δ*P* at *P*_ave_ = 30 kPa (red) or 12.5 kPa (blue). Black: results combining both *P*_ave_ conditions. Inset: CusCBA can dynamically shift between two forms: assembled (stationary, left) and disassembled (mobile, right). OM, outer membrane; PG, peptidoglycan; IM, inner membrane. Yellow star: mEos3.2-tag on CusA. **d**, The diffusion constants of the mobile disassembled state (*D*_m_) and the stationary assembled state (*D*_s_) vs. Δ*P* at different *P*_ave_ or combined. **e**, Copy number of CusA trimers (CusA_3_) at varying Δ*P* and *P*_ave_. **f**, Effective disassembly rate constant *k*_d_ vs. Δ*P* combining both *P*_ave_ conditions. Error bars are s.d. in **c–f**. Numerical values reported here are in mean ± s.d.

When assembled, CusCBA also forms a rigid link across the cell envelope and is therefore subject to mechanical stress and strain in the cell envelope. To understand the effects of mechanical forces on the assembly of CusCBA, we applied extrusion loading to *E. coli* cells (Fig. 1b), in which the inner-membrane protein CusA was tagged by a photoconvertible fluorescent protein mEos3.2 (i.e., CusA^mE^) (Supplementary Note 1.7)^22^. We examined thousands of CusA^mE^ proteins in hundreds of cells and sorted the cells into groups of similar Δ*P* (Fig. 2a). We then used sparse photoconversion and time-lapse stroboscopic fluorescence imaging to track the motions of individual photoconverted CusA^mE^ proteins in each cell at tens of nanometer precision and 60 ms time resolution (Fig. 2b, inset). Within assembled CusCBA complexes the motion of CusA^mE^ is severely restricted but CusA^mE^ that is disassembled from the complex is highly mobile. These two diffusive states of CusA^mE^ can be differentiated by analyzing the distribution of CusA^mE^’s single-molecule displacement lengths (Fig. 2b and Supplementary Note 1.12)^22^. After using deconvolution to correct for cell confinement^22-24^, the displacement length distribution resolves the two diffusive states across all applied pressure conditions: the stationary assembled state and the mobile disassembled state, along with their diffusion constants and fractional populations (Fig. 2b). The resolution and assignment of these two diffusion states were validated previously by control measurements on the free mEos3.2 tag, single deletion strains missing CusC or CusB, and diffusion simulations^22^.

Strikingly, the fractional population of the mobile disassembled state of CusA^mE^ in the cell increases by a factor of ∼2 when Δ*P* increases from ∼6 to ∼33 kPa (Fig. 2c), indicating a direct association between the magnitude of extrusion loading and the disruption of CusCBA assembly in the cell. Concurrently, the effective diffusion constant of the mobile disassembled CusA^mE^ decreases by a factor of ∼4 across this range of Δ*P*, while that of the stationary assembled CusCBA complex, which traverses the cell envelope, remains the same expectedly (Fig. 2d); these trends show that external mechanical stress can influence the CusCBA complex as well as the diffusion of membrane proteins, the latter of which could have contributions from membrane fluidity changes from mechanical stress^25^.

Bacteria submitted to extrusion loading experience a pressure difference across the tapered channels (Δ*P*), as well as a hydrostatic pressure (*P*_ave_, the average between upstream and downstream pressures on a cell), both related to the fluid pressure applied at the device inlet. While the behaviors of CusA^mE^ in response to Δ*P* were substantial, the behaviors of CusA^mE^ showed no significant differences when *P*_ave_ changed by a factor of 2 (Fig. 2c–d, blue vs. red points); combining results from the two *P*_ave_ conditions gave the same behaviors (Fig. 2c–d, black points). Therefore, hydrostatic pressure, at least within our experimental regime, does not play significant roles in membrane protein assembly and diffusivity, suggesting that mechanically-induced disassembly of CusCBA would not be observed using osmotic shock, a commonly used mechanical stimulus that only modifies hydrostatic pressure^26^. Neither CusA^mE^’s copy number nor its spatial distribution shows noticeable changes with varying Δ*P* or *P*_ave_ (Figs. 2e and Extended Data Fig. 5), supporting the idea that modifications in CusCBA assembly induced by extrusion loading are likely not due to cell physiological changes such as CusA protein expression or intracellular localization.

We further analyzed the single-molecule displacement vs. time trajectories of CusA^mE^ to estimate the underlying kinetics of CusA^mE^ disassembly from the CusCBA complex (Extended Data Fig. 6 and Supplementary Note 1.14). The effective disassembly rate constant increases from ∼0.5 to ∼2.8 s^−1^ with increasing Δ*P* (Fig. 2f), supporting the idea that mechanical stress compromises the stability of the assembled CusCBA complex in part by enhancing the disassembly rate.

Sub-lethal concentrations of copper impede the growth of *E. coli*^21^, hence increased disassembly of CusCBA under mechanical stress would suggest a further reduction in cell growth. We thus examined how mechanical stress affected elongation and reproduction of hundreds of individual *E. coli* cells under copper stress by tracking cell length and time to division. Rate of elongation and time to division were both examined in media with 0 or 2.5 mM CuSO_4_ (Extended Data Fig. 7 and 8). The maximum rate of elongation decreased with larger magnitudes of extrusion loading (greater Δ*P*, Fig. 3a) and followed an exponential decay. In the presence of copper stress, the exponential decay rate (0.29 ± 0.11 kPa^−1^, value ± SE) was substantially greater than that without copper stress (0.06 ± 0.03 kPa^−1^), indicating synergy between mechanical and copper stress in suppressing cell elongation (or division). To confirm that the effects of mechanical stress on the function of CusCBA were not limited to extrusion loading in microfluidic chambers, we also assessed the effects of copper stress using an alternate mechanical loading approach: the growth of cells encapsulated in agarose gel with increasing stiffness^27,28^. We used three different concentrations of agarose (0, 0.25, and 0.5 w/v %), corresponding to three different levels of gel stiffness (Extended Data Fig. 9; Supplementary Note 1.13). The maximum growth rate of the *E. coli* population decreased in higher agarose concentration gels, consistent with a previous report^27^ (Fig. 3b, black points). More important, the presence of copper also reduces the growth rate in gels (e.g., pink vs. black points in Fig 3b), and the effect was enhanced in gels with greater agarose concentration. Together, these findings demonstrate that mechanical stress–induced disassembly of CusCBA enhances the toxic effects of copper stress on bacterial physiology.

**Figure 3.**
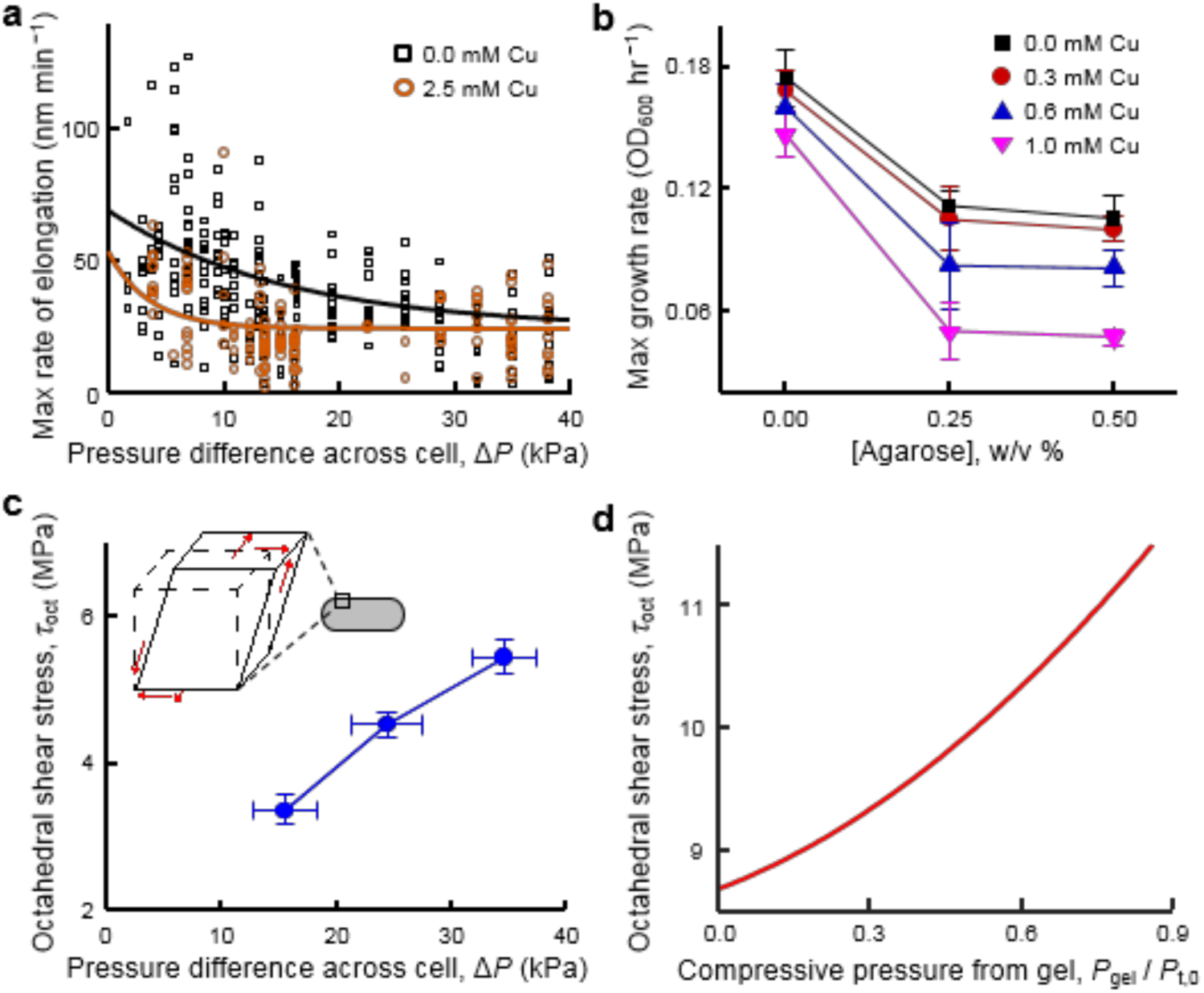
Mechanical loading enhances the toxic effects of copper stress on elongation and growth of *E. coli*. **a**, Maximum rate of elongation of individual cells under extrusion loading without copper stress (*n* = 253 cells) and with copper stress (*n* = 134 cells). Solid lines display exponential decay fits. **b**, Maximum growth rate of cells encapsulated in agarose gel without and with increasing copper stress. Maximum growth rate is influenced by copper concentration, agarose stiffness (concentration) and copper*agarose (p < 0.05), indicating synergy between copper concentration and agarose stiffness. Error bars are s.d. **c**, Finite element analysis demosntrates that octahedral shear stress in the cell envelope of bacteria under extrusion loading increased with increasing Δ*P*. Material properties used in the analysis are in Supplementary Table 3. Error bars are s.d. Inset: 3D depictions of the effects of octahedral shear stress (left) and hydrostatic stress (right) on an infinitesimal element located in the cell envelope on the octahedral plane. Red arrows indicate direction of stresses. **d**, Octahedral shear stress in the cell envelope of bacteria encapsulated in gel (growth confinement loading) increased with increasing compressive pressure from the gel (*P*_gel_). The compressive pressure was normalized to the assumed turgor pressure (*P*_t,0_), where a value of *P*_gel_ / *P*_t,0_ greater than 1.0 would result in buckling or collapse of the cell.

Extrusion loading and gel encapsulation techniques generate substantially different combinations of tensile, compressive, and shear stresses in the bacterial cell envelope (Supplementary Note 1.6). To better understand the components of cell envelope stress associated with mechanically-enhanced disassembly of CusCBA and resulting enhancement of copper sensitivity, we generated analytical and finite element models of the two mechanical loading modalities (Supplementary Notes 1.5 and 1.6). Extrusion loading increases axial tension and reduces tensile hoop stresses while gel encapsulation reduces axial tension in the cell envelope with little effect on hoop stresses (Extended Data Fig. 4c). To identify the forms of mechanical stress that promote disassembly of CusCBA, we decomposed the three-dimensional stress state of the cell envelope into a hydrostatic (volume-changing) component and an octahedral shear (shape-changing) component (Fig. 3c, inset). Hydrostatic stress is known to affect molecular processes like stretch–activated channels, however, in extrusion loading hydrostatic stresses showed a small increasing trend with increasing Δ*P*, whereas in gel encapsulation an opposite trend was observed (Extended Data Fig. 10). The lack of concurrence between the two loading modalities suggests that hydrostatic stress is not likely the main cause of the mechanically-induced disassembly of CusCBA, an assertion that is supported by the fact that CusCBA disassembly during extrusion loading was insensitive to variation in hydrostatic pressure (*P*_ave_, which primarily regulates cell envelope hydrostatic stress). In contrast, mechanical loading through both extrusion loading and gel encapsulation lead to large increases in octahedral shear stress (Fig. 3c–d), suggesting that octahedral shear stress is a likely contributor to disassembly of CusCBA, perhaps by generating forces in the cell envelope that promote movement of the inner and outer membrane components relative to one another and leading to the associated reductions in cell elongation and growth rates (additional discussion in Supplementary Note 1.6).

In summary, our findings demonstrate that mechanical stress in the cell envelope can influence trans-envelope protein complexes resulting in physiologic changes in bacteria. Furthermore, the findings suggest that octahedral shear stresses in the cell envelope such as those generated by adhesion to a surface, overgrowth within crowded cavities, or deformation/growth in small spaces, have the potential to influence the function of other trans-envelope complexes including those involved in cell division, transport of proteins to the outer membrane and resistance to antibiotics. As octahedral shear stress is a well-recognized stimulus for mechanotransduction in mammalian cells and organs^29^, our findings in bacteria suggest a broader role for such stress in cell physiology.

## Supplementary Information

is available in the online version of this paper.

### Acknowledgements

This study was supported by NSF Grant CMMI-1463084 (C.J.H.), NIH Grant GM109993 (P.C.), US Army Research Office W911NF1910121 (P.C.), F31AI143208 (L.A.G.), and a NIH CBI training grant 5T32GM008500 (L.A.G.). This work was performed in part at the CNF, a member of the National Nanotechnology Coordinated Infrastructure (NNCI), which is supported by NSF Grant ECCS-1542081. Imaging on the Andor/Olympus Spinning Disk Confocal at the BRC was supported by NIH Grant 1S10OD010605. We thank the Cornell NanoScale Science and Technology Facility (CNF) and the Biotechnology Resource Center (BRC) staff for assistance, F. Yang for helping with imaging experiments, K. Gunsallus for finite element modeling assistance, and G. Guisado for measurement assistance.

## Author contributions

M.F.R. fabricated microfluidic devices, performed mechanical manipulation, analyzed cell mechanical stresses, and curated data. L.A.G. prepared cell strains, performed single-molecule imaging, analyzed the mechanical effects on efflux complex assembly, and curated data. Y.-C.W. performed mechanical modeling. M.F.R. and L.A.G. performed cell growth assays. A.G.S., B.F., W.J., X.M., C.E.H., L.M.W., and Ł.K. contributed to experiments and data analysis. X.S. contributed to microfluidic device design. A.S. contributed to mechanical modeling. C.-Y.H. supervised mechanical modeling. M.F.R., L.A.G., Y.-C.W., C.E.H., C.J.H., and P.C. analyzed/discussed results and wrote the manuscript. C.J.H. and P.C. conceived and directed research.

## Materials & Correspondence

All data are available from the corresponding authors upon reasonable request.

## Methods

### Fabrication and characterization of microfluidic device

A microfluidic device was used to mechanically stimulate individual bacteria^17^. Six devices were placed onto each fabricated wafer (Extended Data Fig. 1a). The functional units of each device included sixty tapered channels capable of applying extrusion loading at 12 different magnitudes of pressure difference, Δ*P*, and were connected to the device inlet and outlet using feeder channels (Extended Data Fig. 1b-d). Tapered channels were designed with an inlet width large enough to permit entry of individual bacteria (1.2 µm) and exit widths small enough to inhibit bacterial exit (250 nm). Each tapered channel had a length of 75 µm between inlet and outlet. Fluid flow through the bypass channel generated a difference in fluid pressure across each tapered channel.

Sub-micron patterns in the microfluidic devices were fabricated using Deep UV (DUV) lithography at the Cornell Nanoscale Science & Technology Facility. Fused silica was used as a substrate for the microfluidic device to obtain optical clarity and large stiffness. A 55 nm thick chromium etch hard mask was sputter-deposited on silica wafers (500 µm thick, with a 100 mm diameter). A 60 nm thick layer of anti-reflective coating (ARC, DUV 42P) and a 510 nm thick layer of DUV photoresist were deposited on the chromium hard mask using an automated spinner and hot plate system. Exposure of the DUV photoresist was accomplished with a DUV Stepper. The exposed pattern was transferred to the silica substrate using plasma etching. A chrome wet etch was performed to clean off residual chromium. Thru holes were laser-etched into the patterned silica wafer to serve as inlets and outlets.

Depths of device feeder channels were confirmed using a profilometer (Supplementary Fig. 1a). Tapered channel dimensions were confirmed with SEM and AFM (Supplementary Fig. 1b-c). The patterned silica wafer was bonded to a thin, bare silica wafer to seal the device (170 µm thickness). Once bonded, wafers were annealed in a furnace. The wafers were allowed to age for a minimum of one week at room temperature before use.

### Strain construction, sample preparation, and device loading

All strains used in this study were derived from the *E. coli* BW25113 strain (Supplementary Note 1.7). CusA^mE^ was created via lambda Red recombineering, where a short, flexible linker L of 10 amino acids (sequence = AGSAAGSGEF) was used to connect mEos3.2-FLAG (a monomeric, irreversibly photoconvertible fluorescent protein mEos3.2^30,31^ with a C-terminal FLAG tag) to the C-terminus of CusA at its chromosomal locus, as reported in our previous work^22^. This CusA^mE^ fusion protein is functional and stays intact, as previously shown by cell growth assays and Western blot^22^.

To prepare *E.coli* cells expressing mEos3.2-tagged CusA for single-molecule imaging experiments, the cells were first grown in LB medium overnight at 37 °C. The culture was then diluted 1:100 in LB containing chloramphenicol (25 µg/mL) and grown at 37 °C for 4 h (reaching OD_600_ = 0.4). The cells were centrifuged down, washed, and re-suspended in 10 mL of M9 medium supplemented with amino acids, vitamins, and glucose (details in Supplementary Note 1.8). The liquid suspension of cells was loaded into the microfluidic device using syringe pumps and capillary tubing under specific pressures (details in Supplementary Note 1.3).

For cell samples requiring copper stress (e.g., in measuring the rate of cell elongation and division), CusA^mE^ cell cultures were prepared as described above. The pellet was re-suspended in LB, and CuSO_4_ was added to yield a 2.5 mM copper solution. This concentration of copper impacts *E.coli* cell growth but still renders the cells viable.

### Imaging experiments and data analysis

Single-molecule tracking (SMT)^32-38^ via stroboscopic imaging and single-cell quantification of protein concentration (SCQPC) were performed as previously described^22,39^ on an inverted fluorescence microscope (Supplementary Fig. 1 and Supplementary Note 1.9). For SMT, the cells were first illuminated with a 405 nm laser (1–10 W/cm^2^) for 20 ms to photoconvert a single mEos3.2 molecule (or none) from its green fluorescent form to its red fluorescent form. A series of 30 pulses of a 561 nm laser (21.7 kW/cm^2^) in epi-illumination mode with pulse duration *t*_int_ = 4 ms and time lag *T*_tl_ = 60 ms were then used to excite the red mEos3.2. The resulting red mEos3.2 fluorescence was imaged by an EMCCD camera, which was synchronized with the 561 nm laser pulses. This imaging scheme was repeated for 500 cycles for each field of view.

After the SMT step, SCQPC was performed on the same cells, in which the cells were illuminated with the 405 nm laser (1–10 W/cm^2^) for 1 min to photoconvert all remaining mEos3.2 molecules, followed by 561 nm laser illumination for 2000 frames with the same laser power density and exposure time as in the SMT step to quantify the number of remaining mEos3.2 molecules. This 405-illumination and 561-excitation sequence was repeated once more to ensure all mEos3.2 molecules had photobleached. All CusA concentrations cited in the study correspond to those of CusA trimers, i.e., one-third of the total mEos3.2 concentration.

The fluorescence images of individual CusA^mE^ molecules from SMT were analyzed within each fitted cell boundary using a home-written MATLAB program, iQPALM (image-based quantitative photo-activated localization microscopy), the details of which have been reported in our previous study^39^ and described in Supplementary Notes 1.10 and 1.11). Individual CusA^mE^ molecules were identified and position-localized to nanometer precision (typically ∼40 nm precision).

### Agarose embedding assay of cell growth

The procedure for the agarose embedding assay to probe mechanical effects on cell growth was adapted from Auer et al.^28^ The appropriate cells were grown at room temperature in gels of a range of agarose concentrations in cuvettes, which give a range of different mechanical rigidity of gel matrix, and in the presence of a range CuSO_4_ concentrations (details in Supplementary Note 1.13). The cell growth was monitored by absorbance measurements collected every 30 minutes over a period of 5 hours.

### Statistical information

Differences among groups were identified using two-tailed ANOVA. Differences between trends were determined using ANCOVA to account for the effects of covariates. Where appropriate, data were submitted to logarithmic transformation to achieve normal distributions. Least squares regression models were generated to describe trends. Exponential decay rates are determined from nonlinear regression fits to y = a_0_+ a_1_e^(-x\**τ*),^ where *τ* is the decay rate constant (variance noted using SE; e.g., Fig. 3a). Unless otherwise stated, statistical tests were performed with *α* = 0.05. For single-molecule imaging results, the data presented included the number of events observed and the number of protein molecules and cells measured. Standard deviations are provided in relevant figures and tables for data points and fitted parameters.

### Code Availability

Software code is available from the corresponding authors upon reasonable request.

## Extended Data Figures

**Extended Data Figure 1.**
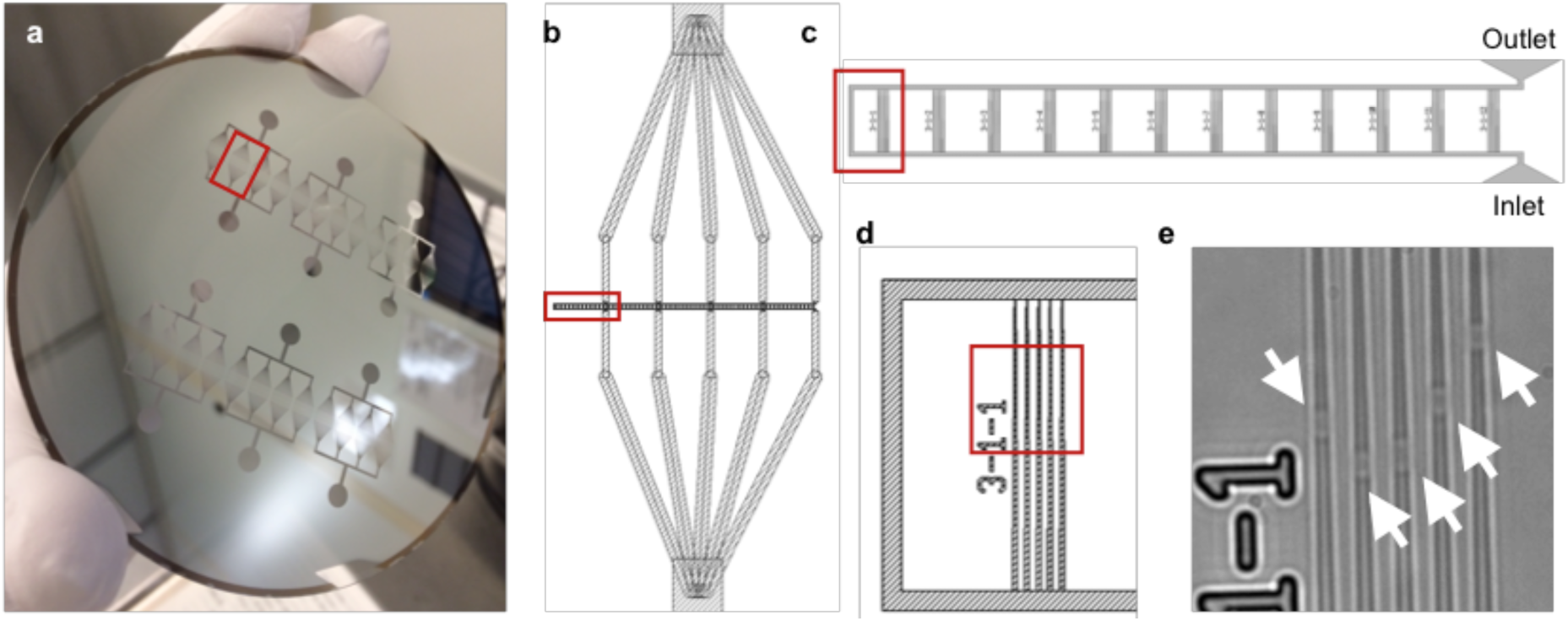
Microfluidic device design. Design and measurements from a typical device are shown. **a**, A patterned wafer shown during the fabrication process, with the chromium hard mask still present. Six devices were patterned into fused silica. Inlets and outlets are seen as circles above and below each device. A red box indicates one subdivision in a device. **b**, Within a subdivision, channels carry bacterial cultures towards five functional units (red box indicating one functional unit). **c**, In one functional unit, twelve groups of five tapered channels are arranged along a bypass channel. **d**, A close-up view of two sets of five tapered channels from the red box in **c** is shown. Numbers adjacent to each set of tapered channels allow for identification and image processing. **e**, A bright-field image of *E. coli* cells (arrows) trapped in five tapered channels.

**Extended Data Figure 2.**
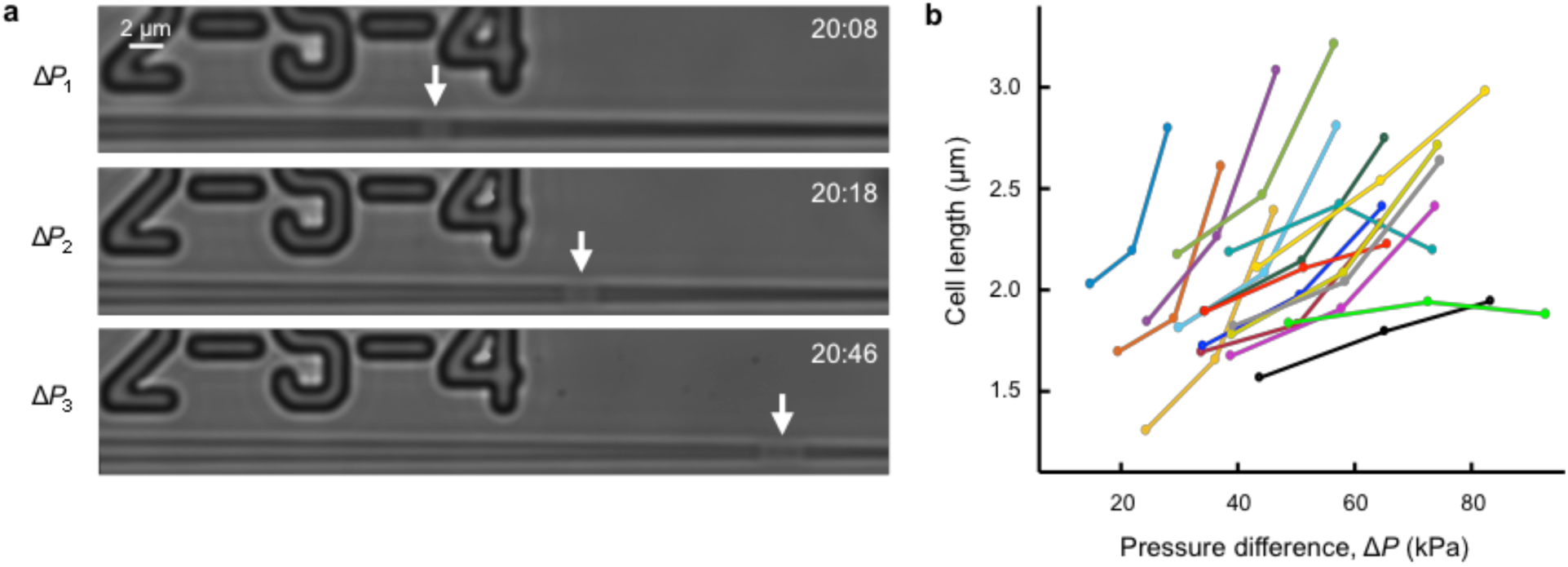
Changes in cell dimensions in *E. coli* submitted to stepwise increases in extrusion loading. **a**, Bright-field images of a trapped cell submitted to three magnitudes of extrusion loading (19.5 kPa, 29.2 kPa and 37.1 kPa top to bottom). Arrow: Trapped *E. coli* cell. **b**, The length of trapped cells at each of the stepwise increases in pressure (*n* = 17; lines connect measures of the same cell).

**Extended Data Figure 3.**
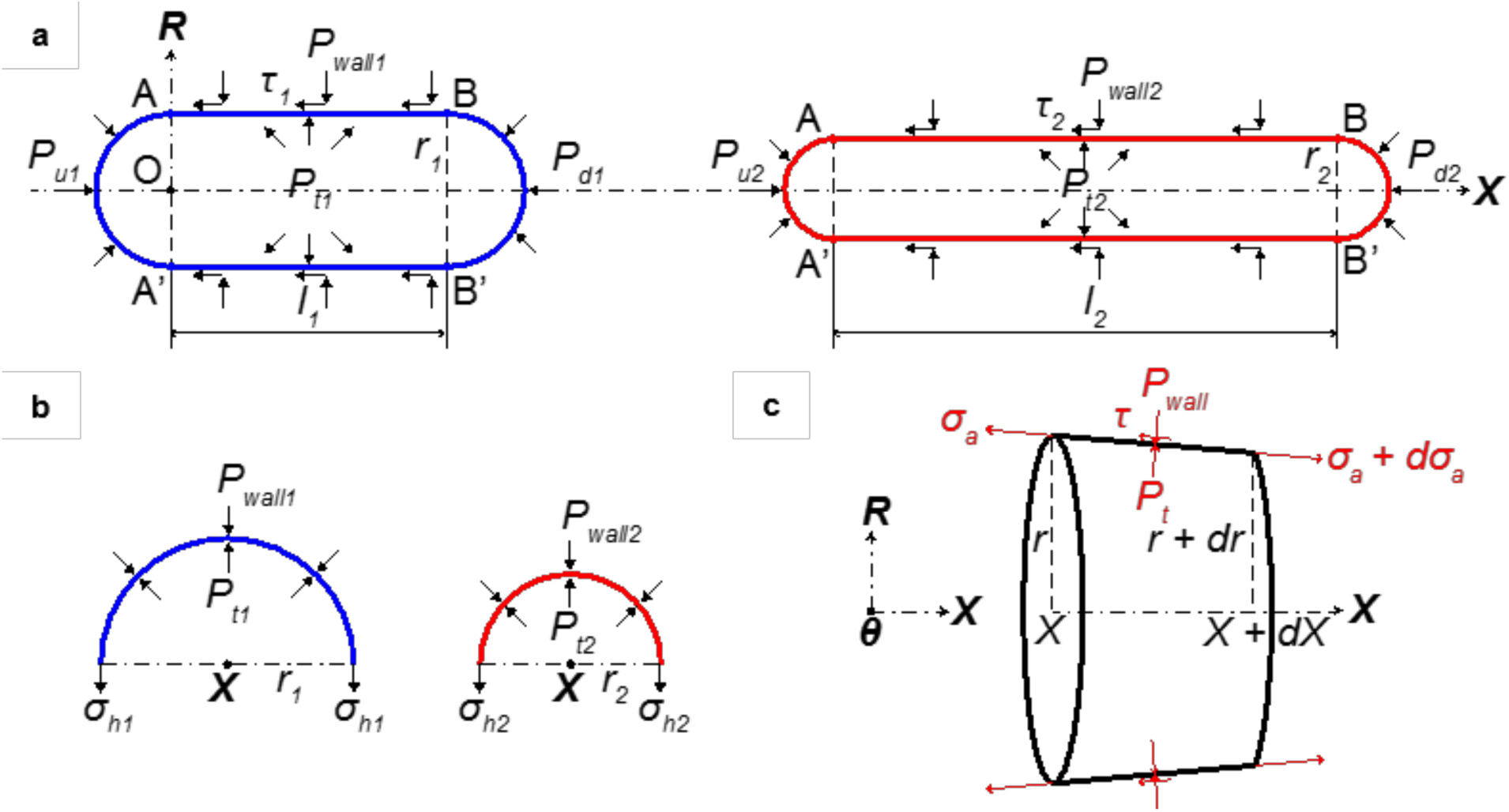
Free body diagrams of a bacterium under extrusion loading. **a**, Schematic of a bacterium trapped inside a channel at two applied pressures. The blue (left) bacterium is at lower applied pressure, and the red (right) one is at higher applied pressure. **b**, The local force balance in the hoop direction. The black dots denote the axial axis. **c**, The local force balance of a segment of a trapped bacterium in the axial direction.

**Extended Data Figure 4.**
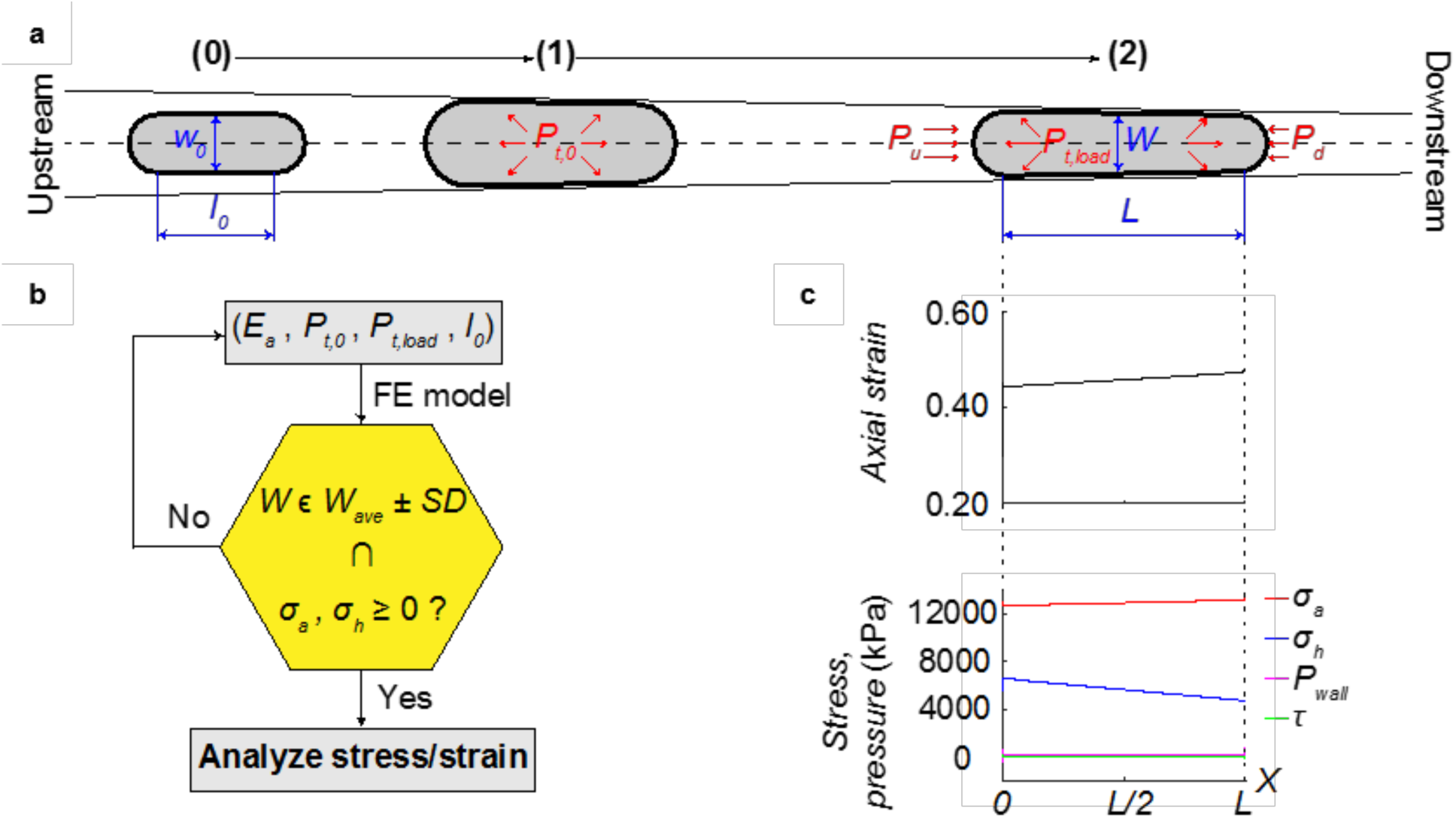
Parametric finite element methods, and the stress and strain results. **a**, Schematic of the finite element model. Numbers mark the steps in a simulation. **b**, Flowchart of the parametric analysis. Finite element model outputs cell width *W*, which is compared to the experimental width plus and minus one standard deviation, *W*_*ave*_ *± SD* (Supplementary Table 2). Axial and hoop stresses are checked whether they are tensile. We analyze the simulative stress and strain states only if these two conditions are achieved. **c**, An example of the simulative stress and strain distributions that match experimental findings. Normal pressure (pink) and axial (red), hoop (blue), and shear (green) stresses are shown for a typical case. Here the variables are: *E*_*a*_ = 25 MPa, *P*_*t,0*_ = 150 kPa, *P*_*t,load*_ = 270 kPa, *l*_*0*_ = 1200 nm, under applied pressure *ΔP* = 20 – 30 kPa, *P*_*ave*_ = 30.0 kPa.

**Extended Data Figure 5.**
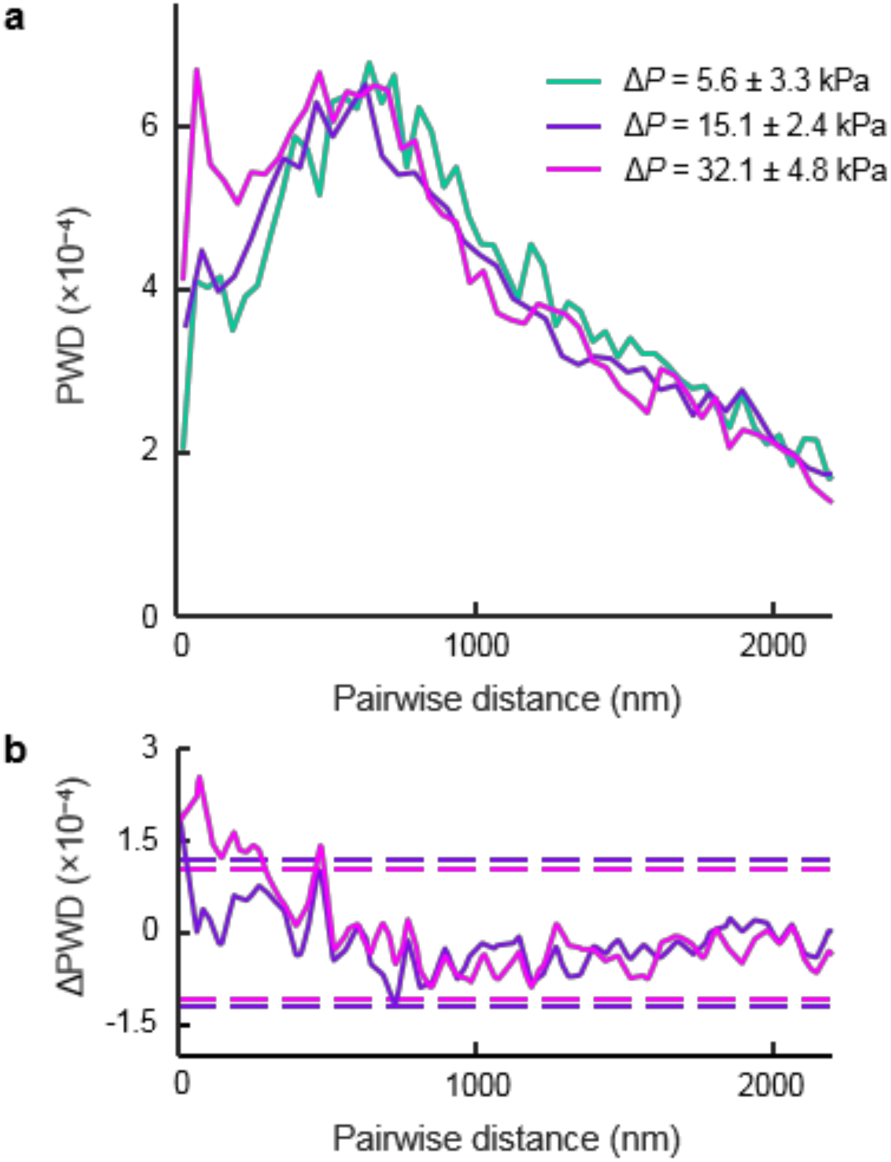
Pairwise distance distribution analysis of CusA^mE^ shows no significant changes under different levels of mechanical stress (i.e., Δ*P*). **a**, Normalized pairwise distance distribution (PWD) obtained from CusA^mE^ SMT results. Green: cells with Δ*P* = 5.6 ± 3.3 kPa; purple: Δ*P* = 15.1 ± 2.4 kPa; pink: Δ*P* = 32.1 ± 4.8 kPa. **b**, Difference of normalized PWDs in A, relative to that of the green curve. The dashed lines are the 99.7% confidence bounds, which represent 3× the standard deviation of the respective ΔPWDs between 2 simulations of randomly-distributed proteins on the cell membrane displaying no clustering: one simulation with 5,000 cells and a total of ∼40,000 to ∼50,000 locations (which give statistically saturated results), and the other with the same number of cells as obtained under experimental conditions (e.g., ∼350 cells in each group with a total of ∼3,000 to ∼3,500 locations), as we previously described^22^. The cell geometry of each simulation matched that of the average cell geometry of the corresponding experimental dataset.

**Extended Data Figure 6.**
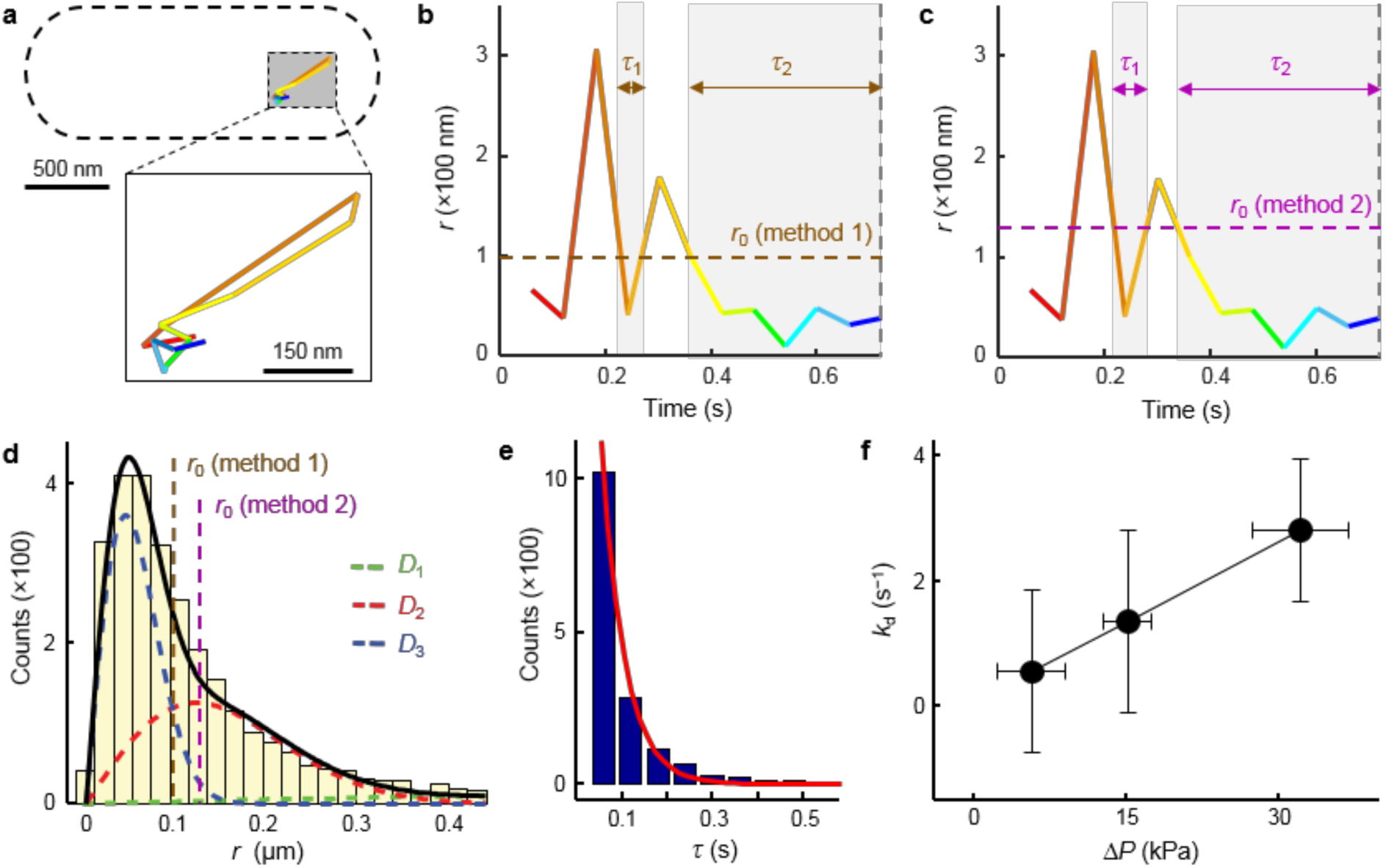
Determination of the effective first-order disassembly rate constant *k*_d_ from SMT trajectories. **a**,**b**,**c**, Position (**a**) and corresponding displacement vs. time trajectories (**b, c**) of a single CusA^mE^ molecule. The horizontal brown dashed line in **b** indicates the threshold *r*_0_ ≈ 100 nm (Method 1), while the horizontal purple dashed line in **c** indicates the threshold *r*_0_ =127 nm (Method 2). Two residence times *τ* are indicated in panels **b** and **c** by the arrows fitting the light gray regions. The vertical gray dashed lines in panels **b** and **c** indicate a photobleaching or photoblinking event. **d**, For Method 1 of determining *r*_0_, the experimental PDF(*r*) before inverse transformation is fitted with three states, with the fastest artificial state in green (*D*_1_) and the two slower states in red (*D*_2_) and blue (*D*_3_), respectively. The overall fit is delineated in solid black. The vertical brown dashed line indicates the threshold *r*_0_ ≈ 100 nm, the intersection between the resolved PDF(*r*) of the 2 slower states. The vertical purple dashed line indicates the threshold *r*_0_ = 127 nm for Method 2. **e**, Distribution of the microscopic residence time *τ* and the fit with Equation S24. **f**, *k*_d_ vs. Δ*P* when using the alternative Method 2, which also shows that *k*_d_ increases with increasing Δ*P*. Error bars are s.d.

**Extended Data Figure 7.**
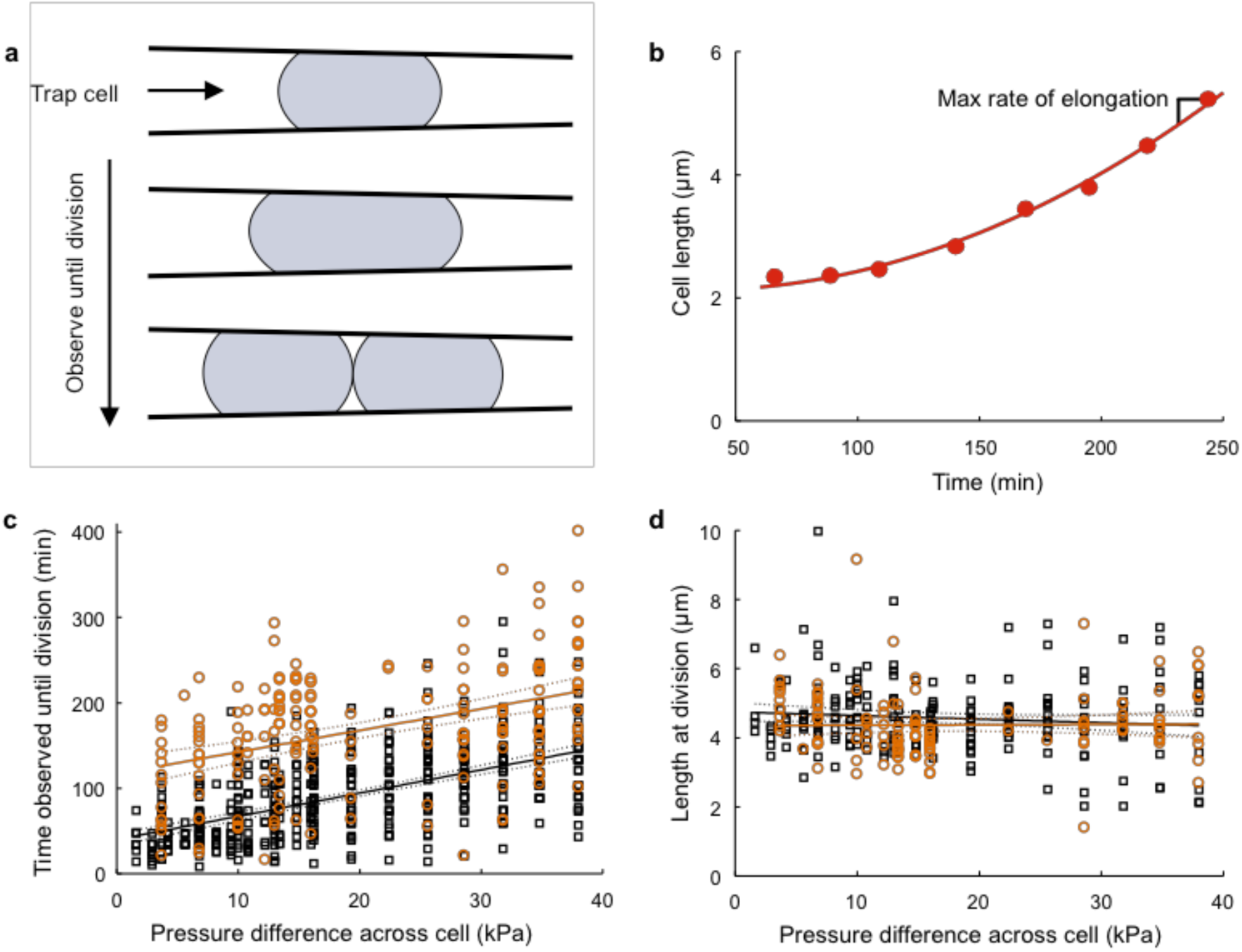
Trapped cells were observed until division, and cellular reproduction was slowed by extrusion loading. **a**, A single cell is trapped at an unknown point during the cell cycle. Sequential images were then taken until the cell was observed to divide. **b**, The cell length was measured using line profiling (see Supplementary Fig. 3) of bright-field images. A quadratic fitting was applied, and the maximum rate or elongation extracted from the fitted line in the time frame was observed. Data from a typical cell is shown. **c**, Time observed until division was plotted for cells not treated with copper (black squares, *n* = 468) and for cells treated with 2.5 mM Cu (orange circles, *n* = 209). An increase in time observed until division was seen for both copper treatment and no treatment cells. All tested pressure difference levels for applied average pressures of 12.5 and 30 kPa saw cell viability. **d**, The length measured at cell division was not noted to change with pressure difference.

**Extended Data Figure 8.**
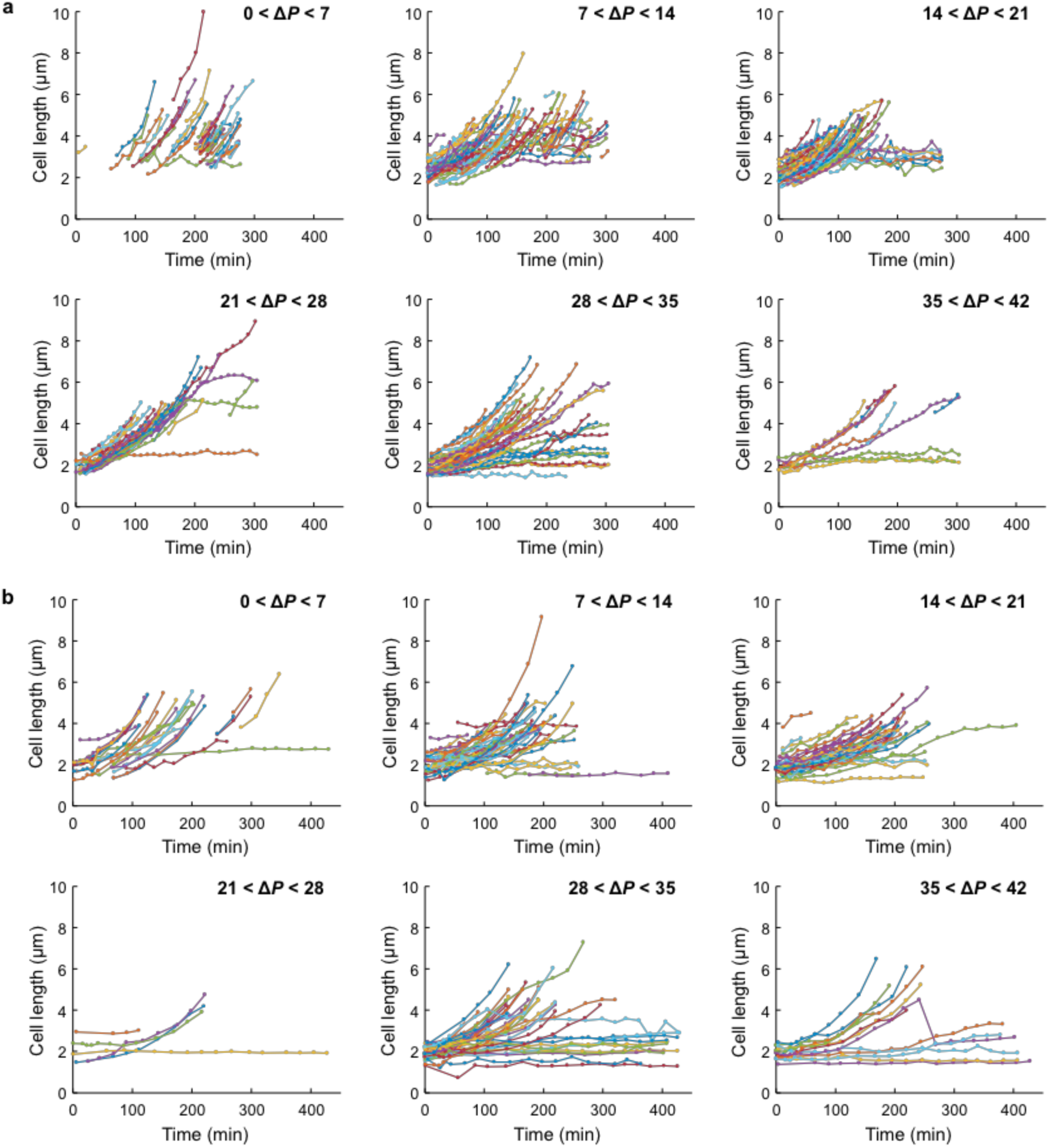
Measured cell length over time for cells that were observed until division. Cells were divided into six pressure difference (Δ*P*) groupings for **a**, no copper treatment (*n* = 260) and **b**, 2.5 mM copper treatment (*n* = 155). Measurements were made as described in Extended Data Fig. 7. Cells where division was not observed are included.

**Extended Data Figure 9.**
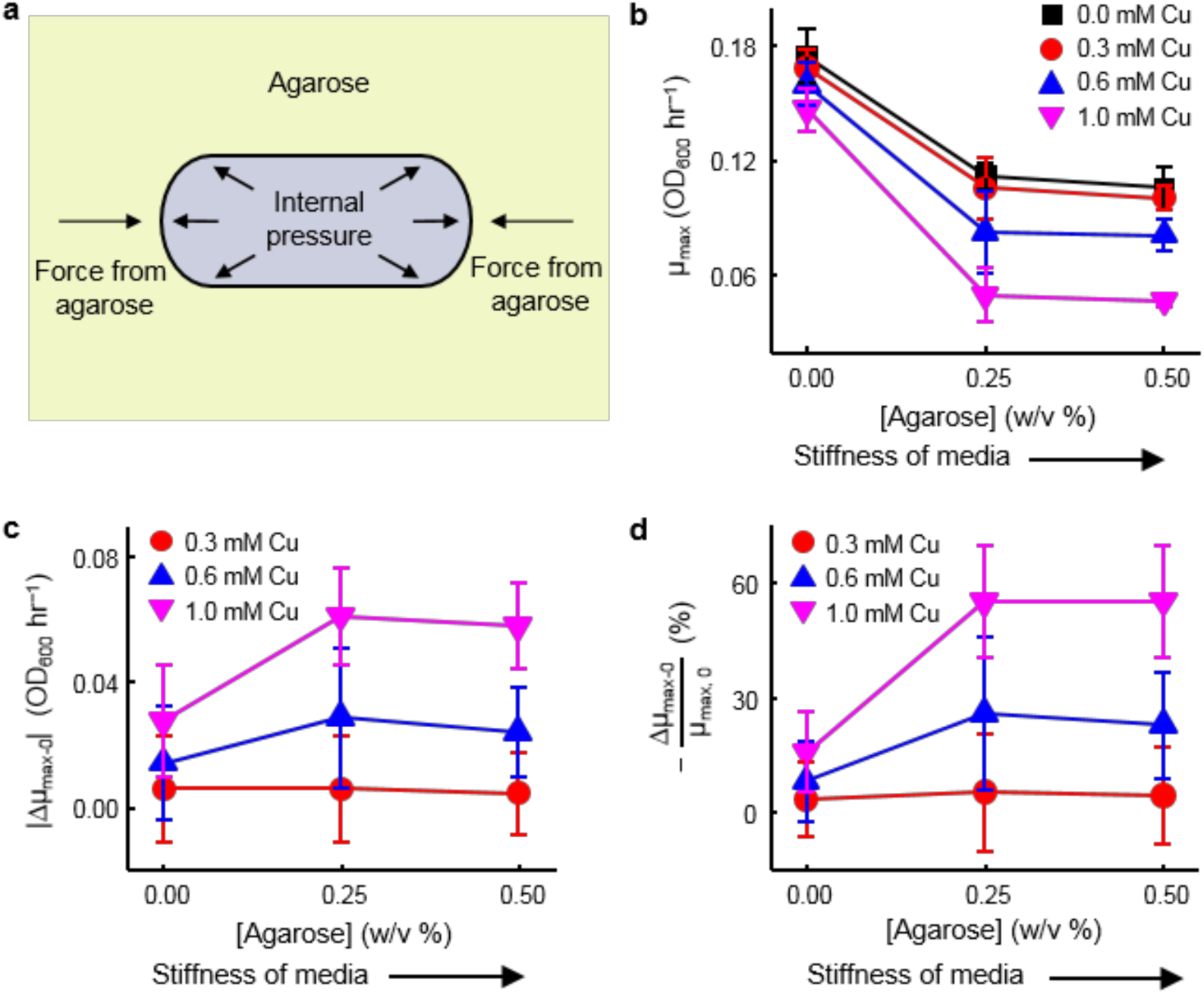
Cells become more sensitive to copper stress at higher agarose concentrations (i.e., more mechanical constraints). **a**, A cell encapsulated in agarose gel is depicted. As the cell grows, the agarose gel exerts a compressive pressure (*P*_cell_) onto the poles of the cell. Internal pressure (*P*_*t*_) within the cell maintains cell shape and prevents buckling. **b**, Maximum growth rate (µ_max_) as a function of percentage of agarose in LB (w/v %). The stiffness of the media increases with increasing agarose concentration. The data in this panel are the same as in Figure 3b in the main text. Maximum growth rate is influenced by copper concentration (p < 0.001), agarose stiffness (concentration, p < 0.001) as well as copper*agarose (p = 0.046), indicating that mechanical stress enhances the effect to copper. Sensitivity of cells to copper stress was determined as the absolute value of change in µ_max_ relative to 0 mM Cu (denoted as µ_max-0_). **d**, Sensitivity of cells to copper stress was determined as the percent decrease in maximum growth rate (µ_max_) relative to 0 mM Cu. Data were recorded in triplicate. Error bars are s.d.

**Extended Data Figure 10.**
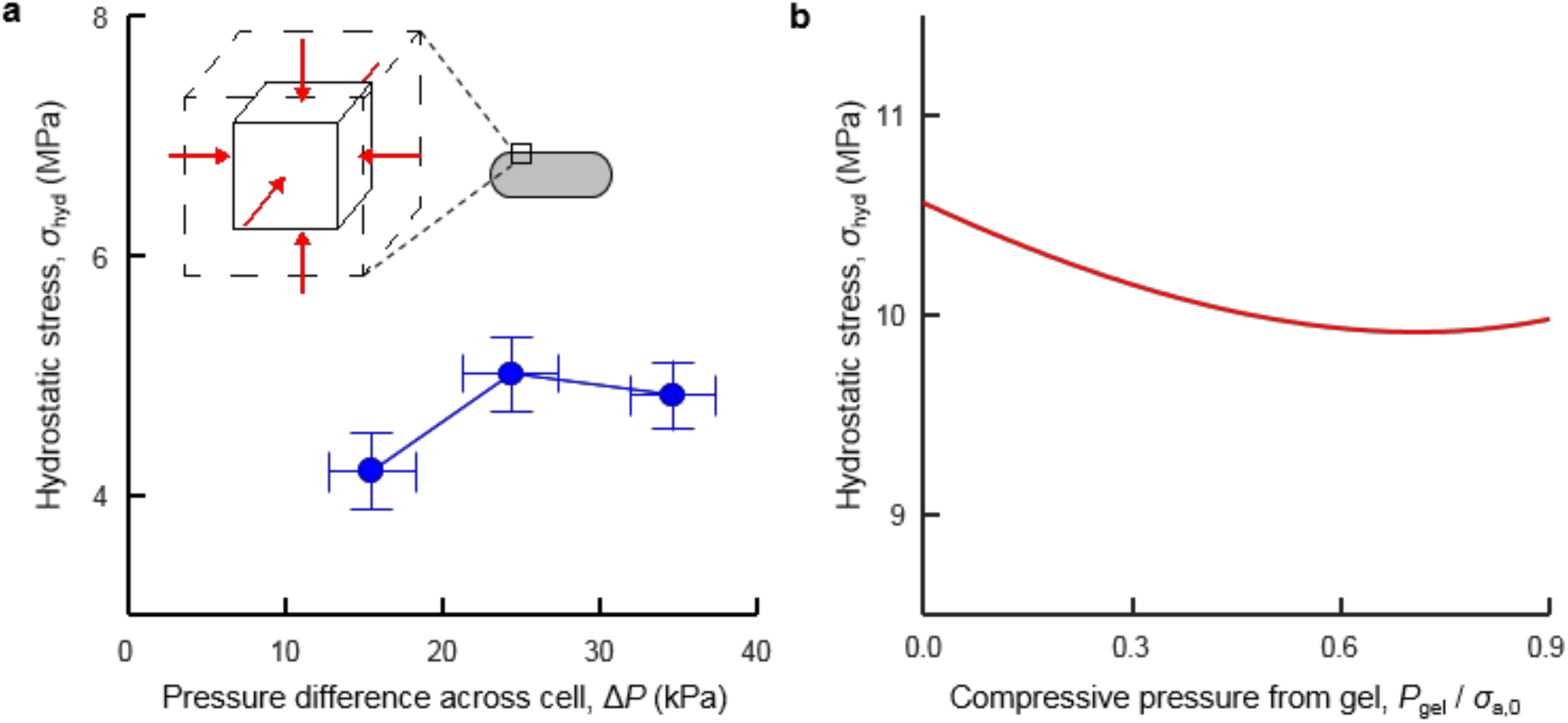
**a.** Hydrostatic stresses (inset) within the cell envelope are not correlated with the magnitude of extrusion loading (ΔP). **b.** Hydrostatic stresses in the cell envelope show only a slight trend with increases in compressive pressure generated by agarose encapsulation.

