## Supplementary Information for "Mechanical stress compromises multicomponent efflux complexes in bacteria"

### 1. Materials and Methods

#### 1.1. Microfluidic device

A microfluidic device was used to mechanically stimulate individual bacteria[1,2]. Six devices were placed onto each fabricated wafer (Extended Data Fig. 1a). Each one of the six devices had four subdivisions, visible as four diamond-like shapes in the device (Extended Data Figs 1a-b). The diamond-like subdivisions contained five fanned-out channels (Extended Data Fig. 1b). Each channel led to a functional unit for extrusion loading experiments (Extended Data Fig. 1b-c). A functional unit consisted of sixty tapered channels arranged in twelve groups of five, with a long bypass channel connecting all tapered channels (Extended Data Fig. 1c-d). Tapered channels were designed with an inlet width large enough to permit entry of individual bacteria (1.2 µm) and exit widths small enough to inhibit bacterial exit (250 nm). Each tapered channel had a length of 75 µm between inlet and outlet. Fluid flow through the bypass channel generated a difference in fluid pressure across each tapered channel. Bacteria submitted to loading within the tapered channels were observed using microscopy (Extended Data Fig. 1e).

Sub-micron patterns in the microfluidic devices were fabricated using Deep UV (DUV) lithography at the Cornell Nanoscale Science & Technology Facility. The device was manufactured from fused silica to achieve optical clarity and large stiffness. Fused silica wafers, 500 µm thick, with a 100 mm diameter (WF3937X02031190, Mark Optics, Santa Ana, CA, USA). Bare wafers were initially cleaned with RCA1 (10 min 6:1:1 $H_2O$:$NH_4OH$:$H_2O_2$ at approximately 70°C) and RCA2 (10 min 6:1:1 $H_2O$:HCl:$H_2O_2$ at approximately 70°C) procedures. A 55 nm thick chromium etch hard mask was sputter-deposited (3 mTorr, 150 W, 450 sec) onto the cleaned surface (Orion 8, AJA International, Scituate, MA, USA). Residue on the bottom of the wafer from the sputter deposition process was removed in a Nanostrip bath (10 min at approximately 50°C). A 60 nm thick layer of anti-reflective coating (ARC, DUV 42P, Brewer Science, Rolla, MO, USA) and a 510 nm thick layer of DUV photoresist (UV210, MicroChem, Westborough, MA, USA) were deposited on the chromium hard mask using an automated spinner and hot plate system (Gamma, Suss MicroTec AG, Garching, Germany). Thickness and uniformity of the ARC and DUV photoresist layers were validated by optical measurement (FilMetrics, San Diego, CA, USA). Exposure of the DUV photoresist was accomplished with a DUV Stepper at approximately 28 mJ/cm$^2$ exposure and approximately −0.14 µm focus offset, 0.63 numerical aperture, and 0.8 outer sigma (ASML 300C, Veldhoven, Netherlands). The exposed photoresist was then developed for 60 sec with a single puddle (AZ 726 MIF, MicroChemicals GmbH, Ulm, Germany).

The exposed pattern was transferred to the silica substrate using plasma etching. An Ar/$O_2$ etch was performed to transfer the pattern from the DUV photoresist to the ARC layer (1.5 min, 42.5 sccm Ar, 7.5 sccm $O_2$, 15 mTorr, PlasmaLab 80+ , Oxford Instruments, Oxfordshire, England), followed by a $Cl_2$/$O_2$/Ar etch to transfer the pattern to the chromium hard mask below (90s, 1 sccm $Cl_2$, 9 sccm $O_2$, 3 sccm Ar, 30 mTorr, Minilock III, Trion Technology, Tempe, AZ, USA). Oxygen plasma was used to remove residual photoresist and ARC (3 min, 50 sccm $O_2$, 60 mTorr, Oxford 80+). The exposed chromium pattern was then transferred to the silica substrate using a $CHF_3$/Ar etch (4:35 min, 30 sccm Ar, 20 sccm $CHF_3$, 4 mTorr, 50°C). A chrome wet etch was performed to clean off residual chromium (Cyantek CR-14, KMG, Fort Worth, TX,

USA), leaving behind bare silica with the etched device pattern. Thru holes were laser-etched into the patterned silica wafer to serve as inlets and outlets (VersaLaser VLS3.50, Universal Laser Systems, Scottsdale, AZ, USA). Ridges in the silica resulting from the laser etch were removed with a 10 sec 6:1 BOE dip coupled with abrasions from a diamond drill bit (1 mm diamond drill, Lasco Diamond Products, Chatsworth, CA, USA).

Characterization was performed to confirm channel depths (Supplementary Fig. 1a). Depths of device feeder channels were confirmed using a profilometer (P10, KLA-Tencor, Milpitas, CA, USA). Characterization of taper dimensions was performed with SEM (UltraSEM, Zeiss, Oberkochen, Germany) and AFM (Icon, Veeco, Plainview, NY, USA), where taper inlet and outlet widths were measured with SEM images and channel depths were sampled with a high-aspect ratio AFM tip (Supplementary Fig. 1b-c). Once device characterization was finished, the patterned silica wafer was bonded to a thin, bare silica wafer to seal the device (170 µm thickness, WF3937X02031190, Mark Optics). A wafer thickness of 170 µm is equivalent to a no.1 thickness microscope slide. The fused silica wafers were put through a MOS clean sequence of RCA1, RCA2, and RCA1 baths before drying in a spin-rinse-dryer. Immediately afterwards, the patterned and bare silica wafers were bonded together and annealed in a furnace tube at 1100°C for 5 hr in a $N_2$ environment. Once bonded, wafers were removed from the furnace and allowed to sit for a minimum of one week at room temperature before use.

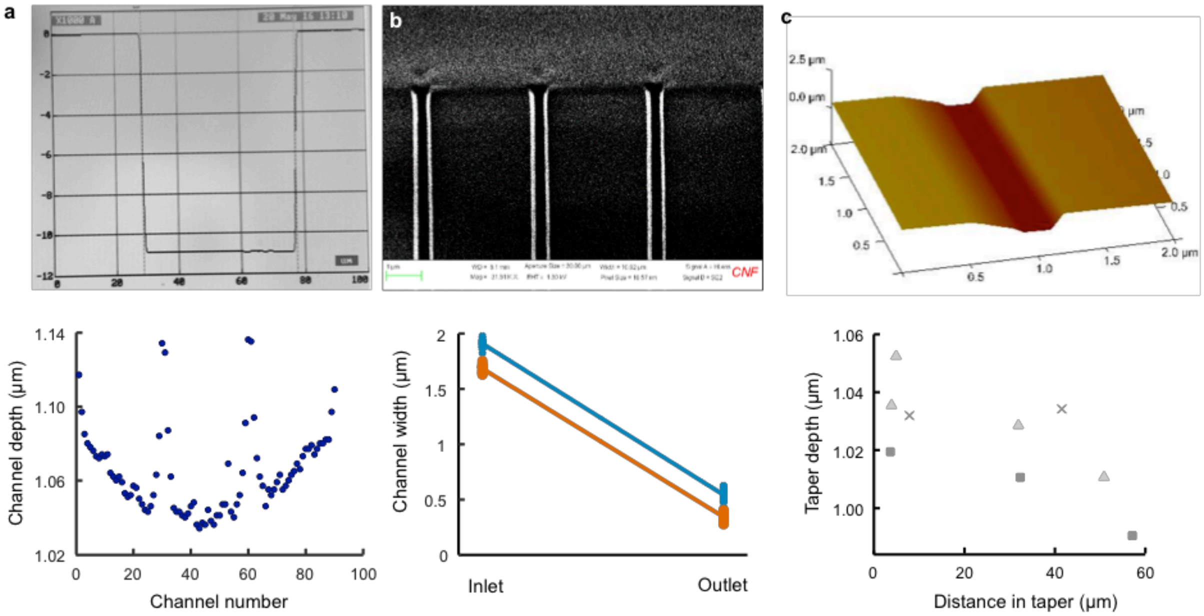

**Supplementary Figure 1.** Validation of microfluidic dimensions was performed using profilometry, SEM, and AFM. Measurements from a typical device are shown. **a**, Prior to sealing devices, fluidic channels with large widths were scanned by a profilometer. Variations in channel depth within the testing region of the device were no more than 10%. **b**, Inlet and outlet widths for tapered channels were measured by SEM. Variation between wafers could be large (up to 60%), but variation within one experimental device was smaller (20%). **c**, AFM was used to verify tapered channel depths were less than 1200 nm. Sample measurements from three wafers are shown and indicated using distinct symbols.

#### 1.2. Fluid pressure calculation

The fluidic pressure through the device was determined from the inlet pressure (measured) and hydraulic circuit model[3]. Fluidic pressure was determined using the Hagen–Poiseuille law (Eq. S1), where $Q$ is the flow rate, $\Delta P$ is the pressure difference between the inlet and outlet of a channel, and $R_h$ is the hydraulic resistance of the channel.

$$\Delta P = QR_h \qquad \text{Eq. S1}$$

Poiseuille flow was assumed for channels where the ratio between width and depth was small (width/depth < 20). The hydraulic resistance determined using Poiseuille flow is shown in Eq. S2, where $\mu$ is the fluid viscosity, $L$ is the channel length, $A$ is the cross-sectional area, and $r_h$ is the hydraulic radius of the channel[3].

$$R_h = \frac{8\mu L}{Ar_h^2} \qquad \text{Eq. S2}$$

The hydraulic radius is an equivalent length of the channel's geometry, defined below in Eq. S3 where $P$ is the channel perimeter.

$$r_h = \frac{2A}{P} \qquad \text{Eq. S3}$$

Plane Poiseuille flow was assumed when the ratio between width and depth of the fluidic channel was large (width/depth > 20). Hydraulic resistance determined for plane Poiseuille flow is given in Eq. S4, where $h$ is the height (depth) of the channel.

$$R_h = \frac{12\mu L}{Ah^2} \qquad \text{Eq. S4}$$

#### 1.3. Loading of device – protocol

Fluid was delivered to the device using a syringe pump (Fusion Infusion 400 Syringe Pump, Chemyx, Stafford, TX, USA) with a 3mL syringe (Monojet, Medtronic, Minneaplis, MN, USA) connected to capillary tubing (OD 360 µm, IDEX-HS 1572, IDEX, Lake Forest, IL, USA) (Supplementary Fig. 2). The capillary tubing and syringe were joined together with a luer adapter (P-642, IDEX) and union assembly (VICI ZRU1.5XC union, VICI C-NL.35L sleeve, VICI C-NNFFPK ferrule extracted from fitting, VICI Valco Instruments, Houston, TX, USA). A pressure sensor (uPS0800-C360-10, LabSmith, Livermore, CA, USA) was attached to the capillary tubing line using an interconnect tee (C360-203, LabSmith) and threaded fittings (C360-100, LabSmith). A seal was formed between the capillary tubing and device inlet using a fluidic attachment (BMP-LP-MAG, CorSolutions, Ithaca, NY, USA) and flat-bottomed PEEK ferrule (N-123-03, IDEX).

A solution of 70% EtOH (0.5 mL) was flushed through the capillary tubing to clean out any residue, followed by sterile growth media to prime the tubing line. For super-resolution microscopy, 1X-M9 minimal media was used (see "1.8. Cell sample preparation") as the growth media. For diffraction-limited microscopy, sterile LB Broth was used (see "1.8. Cell sample preparation"). For copper-treated media, sterile LB Broth was treated with 2.5 mM of $CuSO_4$ (see "1.8. Cell sample preparation").

After flushing the lines with experimental media, the capillary tubing was attached to the device and connected with the fluidic attachment. Force was applied on a syringe plunger to increase fluidic pressure. The pressure in the fluid was measured and maintained at approximately 80 kPa for 6 min to wet the channels and flush out air bubbles. Once the device channels were wetted, the tubing line was disconnected from the device. The capillary tubing was then flushed with 0.25 mL of media with bacteria.

To load bacteria into the device, the syringe with bacteria was removed and exchanged for a syringe with sterile media to prevent bacterial concentrations from becoming too great. The capillary tubing was then reconnected to the device with the fluidic attachment. The media-filled syringe connected to the capillary tubing was mounted onto a syringe pump, and pressure was increased until bacteria were observed within the tapered channels (Fusion 400 Syringe Pump, Chemyx, Stafford, TX, USA). Once bacteria were loaded into the tapered channels, the pressure was reduced to the desired experimental value (the magnitude of pressure while loading was not found to influence results).

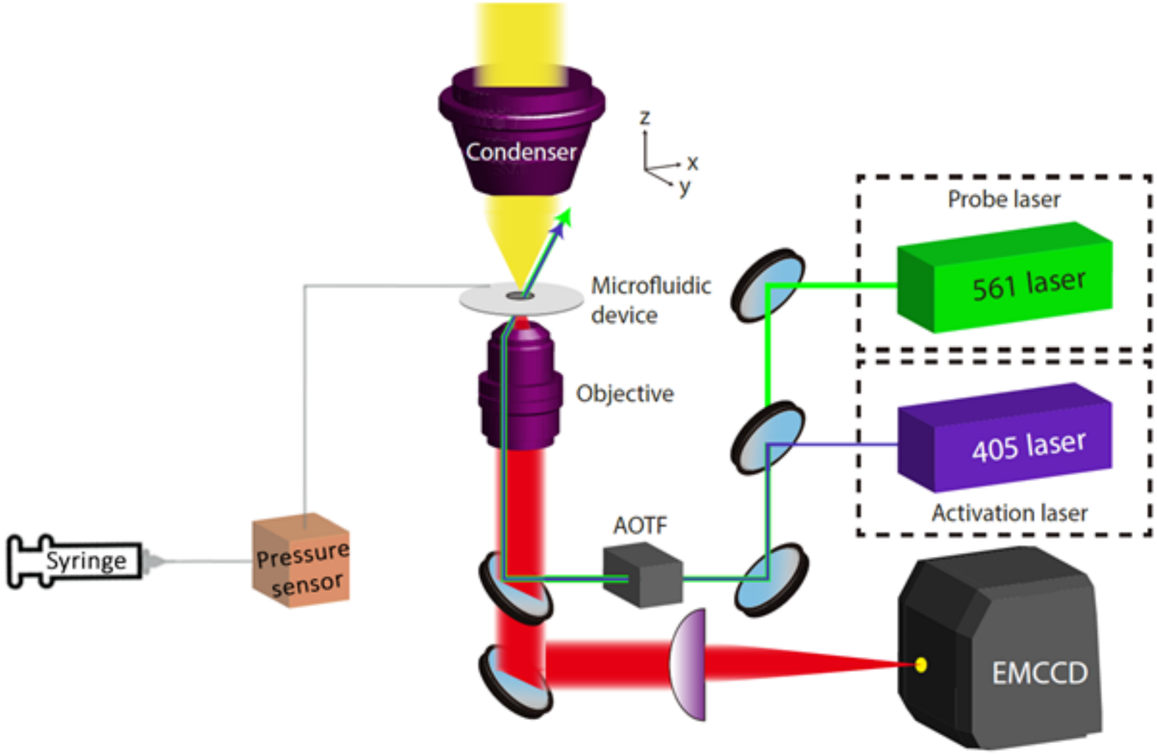

**Supplementary Figure 2.** Extrusion loading implemented with previously demonstrated PALM imaging[2,4]. A syringe pump was used to apply fluidic pressure. Pressure was measured along the capillary tubing line prior to entering the microfluidic device.

#### 1.4. Deformations of bacteria submitted to stepwise increases in extrusion loading

*E. coli* suspended in liquid media were submitted to extrusion loading (Fig. 1a-b) with an initial $P_{ave}$ of 46 kPa ($\Delta P$: 15–49 kPa). Microscopy images of trapped bacteria were collected using bright-field microscopy (0.108 µm per pixel, Extended Data Fig. 1e). To induce a stepwise increase in extrusion loading on trapped cells, the applied fluidic pressure was increased (measured at the device inlet). The trapped cells were imaged at the larger applied pressure again using bright-field microscopy (Extended Data Fig. 2a). The process was repeated for another stepwise increase in applied pressure, resulting in an image of the same trapped cells at three $\Delta P$ magnitudes. Stepwise loading occurred over a short timeframe (~19 min) at room temperature to minimize the influence of cell growth on changes in cell length. Increases in applied $\Delta P$ resulted in movement of bacteria farther into the tapered channels leading to a reduction in cell width (Extended Data Fig. 2a-b). Additionally, cell length increased (Extended Data Fig. 2b). These observations show that increased magnitude of extrusion loading results in reductions in tensile hoop strains, increases in tensile strain in the axial direction, and a net reduction in cell volume (Fig. 1c-d).

#### 1.5. Analytical model of cell envelope stress state

An analytical model was used to characterize the stress and strain states in an *E. coli* cell submitted to stepwise increases in extrusion loading. The model uses the thin-walled approximation and assumes an axisymmetric cell geometry. The cell contains a turgor pressure ($P_t$) associated with osmolarity and is submitted to an upstream hydrostatic pressure ($P_u$) and a downstream hydrostatic pressure ($P_d$). The difference between upstream and downstream pressure ($P_u - P_d$) is the pressure difference $\Delta P$. A normal pressure ($P_{wall}$) is exerted by the channel wall on the cell, and a frictional stress ($\tau$) associated with contact to the cell envelope is also present (Extended Data Fig. 3a).

The stress and strain, and the change in turgor pressure of a bacterium at two loading configurations ($\Delta P_1$ and $\Delta P_2$), are considered (Extended Data Fig. 3a). The cylindrical portion of the bacterium (AA' to BB' in Extended Data Fig. 3a) is referred to as 'trunk' and the ends caps as 'caps.' The cell envelope thickness is denoted by $t$. As the taper angle $\alpha$ in the microfluidic channel is very small (~0.5°), we assume uniform radius for the trunk.

We denote the displacement of the point $X$ in the cell envelope as it moves from configuration 1 to configuration 2 as $u$, such that $x = X + u$. As the cell moves from configuration 1 to configuration 2, the axial strain $\varepsilon_a$ and the hoop strain $\varepsilon_h$ in the cell envelope are given by:

$$\varepsilon_a = du(X)/dX = u'(X) \qquad \text{Eq. S5}$$

$$\varepsilon_h = \frac{r_2 - r_1}{r_1} \qquad \text{Eq. S6}$$

We assume small strains and neglect bending of the cell envelope[5]. The incremental tensile axial stress $\delta\sigma_a$ and the incremental tensile hoop stress $\delta\sigma_h$ between two configurations are given as:

$$\sigma_{a2} = \sigma_{a1} + \delta\sigma_a \qquad \text{Eq. S7}$$

$$\sigma_{h2} = \sigma_{h1} + \delta\sigma_h \qquad \text{Eq. S8}$$

The cell envelope is modeled as a linear elastic transversely isotropic material[6,7] with hoop Young's modulus, $E_h$, on the transverse isotropic plane and the axial Young's modulus, $E_a$. The ratio between hoop and axial Young's modulus is described by the anisotropic coefficient $\gamma = E_h / E_a$. The Poisson's ratio in the axial direction is denoted by $\nu_{ha}$. The transversely isotropic constitutive law between incremental stresses ($\delta\sigma_a$ and $\delta\sigma_h$) and strains ($\varepsilon_a$ and $\varepsilon_h$) gives the following equations:

$$\delta\sigma_a = \frac{E_h}{\gamma - \nu_{ha}^2}(\varepsilon_a + \nu_{ha}\varepsilon_h) = \frac{E_h}{\gamma - \nu_{ha}^2}\left[u'(X) + \nu_{ha}\frac{r_2 - r_1}{r_1}\right] \qquad \text{Eq. S9}$$

$$\delta\sigma_h = \frac{E_h}{\gamma - \nu_{ha}^2}(\nu_{ha}\varepsilon_a + \gamma\varepsilon_h) = \frac{E_h}{\gamma - \nu_{ha}^2}\left[\nu_{ha}u'(X) + \gamma\frac{r_2 - r_1}{r_1}\right] \qquad \text{Eq. S10}$$

Local force balance in the hoop direction (Extended Data Fig. 3b) for the two pressure configurations can be expressed as:

$$\sigma_{h1}(X) = \frac{r_1}{t}\left[P_{t1} - P_{wall1}(X)\right] \qquad \text{Eq. S11}$$

$$\sigma_{h2}(X) = \frac{r_2}{t}\left[P_{t2} - P_{wall2}(X)\right] \qquad \text{Eq. S12}$$

Axial force balance (Extended Data Fig. 3c) is obtained by assuming a constant Coulomb friction coefficient $f$ between the cell and channel walls, such that the shear friction stress was given by $\tau = f \cdot P_{wall}(X)$. The resulting axial force balances for both configurations are given in Eqs. S13 and S14, with higher-order incremental terms not included.

$$\frac{d\sigma_{a1}(X)}{dX} = \frac{f}{t}P_{wall1}(X) + \frac{\alpha}{r_1}\left[\sigma_{a1}(X) - \sigma_{h1}(X)\right] \qquad \text{Eq. S13}$$

$$\frac{d\sigma_{a2}(X)}{dX} = \left\{\frac{f}{t}P_{wall2}(X) + \frac{\alpha}{r_2}\left[\sigma_{a2}(X) - \sigma_{h2}(X)\right]\right\}(1 + u') \qquad \text{Eq. S14}$$

Considering the cell as a single body, the axial forces from the pressure difference must be balanced by wall pressure and friction, which results in the following equations:

$$\hat{P}_{wall1} = \frac{1}{l_1}\int_0^{l_1} P_{wall1}(X)dX = \frac{r_1 \Delta P_1}{2l_1(f+\alpha)} \qquad \text{Eq. S15}$$

$$\hat{P}_{wall2} = \frac{1}{l_2}\int_0^{l_1} P_{wall2}(X)(1+u')dX = \frac{r_2 \Delta P_2}{2l_2(f+\alpha)} \qquad \text{Eq. S16}$$

The change in turgor pressure between the two configurations can be assessed from the force balance in the hoop direction. By subtracting Eq. S11 from Eq. S12, then integrating both sides of the equation across the length of the cell and using Eqs. S10, S15, and S16, we obtain:

$$P_{t2} - P_{t1} = \frac{r_1(\Delta P_2 - \Delta P_1)}{2l_1(f+\alpha)} + \frac{tE_h}{r_1(\gamma - \nu_{ha}{}^2)}\left(\nu_{ha}\frac{l_2 - l_1}{l_1} + \gamma\frac{r_2 - r_1}{r_1}\right) \qquad \text{Eq. S17}$$

Substituting values for pressure, radius, and Young's moduli within the experimental range, the right-hand side of Eq. S17 is always positive (i.e. increases in pressure difference result in increases in internal pressure). The increase in turgor pressure is consistent with observed reductions in cell volume during stepwise extrusion loading (Fig. 1C). Since the cell remains viable after loading, it is expected that the volume loss is predominately water. A reduction in water content would be expected to increase osmolarity and thereby increase turgor pressure.

Differentiating Eq. S7 and replacing the resulting terms with Eqs. S9, S11-14, and S17 gives us Eq. S18, where *M* is given by Eq. S19 and *B* by Eq. S20.

$$u'' + Bu' = M \qquad \text{Eq. S18}$$

$$M \triangleq \frac{\left(\gamma - \nu_{ha}^2\right)f}{tE_h}\left[\frac{r_1\left(\Delta P_2 - \Delta P_1\right)}{2l_1\left(f+\alpha\right)} + \frac{tE_h\nu_{ha}\frac{l_2 - l_1}{l_1}}{r_1\left(\gamma - \nu_{ha}^2\right)}\right] + \frac{\alpha\left(\nu_{ha} - \gamma\right)\varepsilon_h}{r_1} \qquad \text{Eq. S19}$$

$$B \triangleq \frac{(f+\alpha)\nu_{ha} - \alpha}{r_1} \qquad \text{Eq. S20}$$

The displacement ($u$) and the axial strain ($\varepsilon_a$) distribution can be obtained by solving (S18) while noting that cell geometry requires [$r_2 - r_1 = -u(0) \cdot \alpha$] and [$u(l_1) - u(0) = l_2 - l_1$]. The axial strain distribution is found to be:

$$\varepsilon_x = u'(X) = \frac{M}{B} - \frac{Ml_1 - B\left(l_2 - l_1\right)}{1 - e^{-Bl_1}} e^{-BX} \quad \text{Eq. S21}$$

The axial strain can be shown to increase monotonically from the upstream end ($X = 0$) to the downstream end ($X = l_1$) of the trunk if we solve and use the results of the force balance at two end caps in the axial direction. The incremental axial stress is then obtained from Eqs. S9 and S21, which also exhibits a monotonically-increasing pattern from the upstream end to the downstream end of the cell trunk.

#### 1.6. Finite element model for bacteria submitted to mechanical loading

The experimental results (Fig. 3c-d) indicate a response to both extrusion loading and loading through gel encapsulation. The analytical model described above, while providing insight into the stresses and strains generated by extrusion loading, has limited utility for comparing the two loading modes since many of the variables are not well characterized. Here, we generate finite element models to compare cell envelope stress states between the two loading modes.

Characterization of the stress and strain states in the cell envelope during extrusion loading was performed using nonlinear finite element analysis. A finite element model of a bacterium submitted to extrusion loading was developed using Abaqus (CAE 6.9-EF2, Dassault Systems, Providence, RI, USA). An axisymmetric model was generated in which the cell envelope consisted of solid elements. The use of solid elements constrains the model to consider situations with a positive turgor pressure (a reasonable assumption given the increases in internal pressure during extrusion loading, see above). The trunk of the cell envelope was modeled as transversely isotropic (see above section). The Poisson's ratio was modeled as 0.3 on the isotropic plane and as 0.34 in the hoop/axial direction[8]. The cell end caps were assigned isotropy, with the Young's modulus set as the average of the Young's moduli in the hoop and axial direction of the trunk. The Poisson's ratio in the end caps was assumed as 0.3. Channel walls were represented as rigid surfaces with dimensions matched to the microfluidic device design. Contact between the cell envelope and the channel walls was simulated using a surface-to-surface contact. Coulomb friction was applied between the cell envelope and channel walls. Each finite element model consisted of 103,680 four-node bilinear quadrilateral elements (16 elements across the cell thickness, CAX4 elements with geometric nonlinearities included). The thickness of the cell envelope $t$ was assumed to be 4 nm[6,9].

Extrusion loading was simulated in two steps (Extended Data Fig. 4a). In the initial step, the cell envelope is inflated with a turgor pressure characteristic of the free-floating state ($P_{t,0}$). Although reports of turgor pressure vary dramatically in the literature, cell width is highly constrained[10] and has small variability in the strain of *E. coli* used in this experiment[2]. Hence, we expect a relationship between unstressed width $w_0$, turgor pressure, and the Young's modulus of the cell envelope. The unstressed width at a given turgor pressure and cell envelope Young's modulus were determined iteratively to achieve a final width that was the same as the cell width of free-floating *E. coli*.

In the second step of the simulation, upstream and downstream pressure were included and the internal pressure was increased to $P_{t,load}$. Stabilization control was used to allow for small rigid body motion of the cell envelope within the channels and was confirmed to not affect final stress and strain (data not shown).

As many of the parameters used in the finite element model have not yet been well defined experimentally (Supplementary Table Supplementary Table 1), we performed a parametric analysis to identify the combination of descriptive variables that result in deformations similar to what is seen experimentally (Extended Data Fig. 4b). Key input parameters for the model include Young's moduli, initial turgor pressure, and turgor pressure after deformation. Finite element model results were compared to experimentally-measured cell widths and lengths with a known pressure difference $\Delta P$ (data from Supplementary Table Supplementary Table 2). Cell width was not highly variable in this strain of *E. coli*, but cell length varied considerably depending on the cell cycle. For this reason, final deformed cell width was used as the primary indicator of a finite element model result consistent with experimental findings. If the deformed cell width was within one standard deviation of the experimental cell width, the finite element model result was accepted. The parametric analysis was performed with the following parameter values: the Young's modulus in the hoop direction $E_h$ was two times larger than the Young's modulus in the axial direction $E_a$[6,7]. The friction coefficient $f$ was estimated from the global force balance on the cell envelope in the axial direction of a trapped cell. When we ran simulations with $P_{t,0}$ as 100, 150, and 200 kPa, $f$ was estimated to be 0.0039, 0.0026, and 0.0020, respectively. The unstressed cell length $l_0$ can also affect how far a cell travelled downstream into the tapered channel, affecting the final cell width, so $l_0$ was parameterized as well. Although the amount of initial contact between the cell envelope and the channel walls in the first step of the simulation can vary based on the axial position of the cell within the tapered channel, variation in the starting position in the channel had little effect on final cell width. A change of 78% in the starting position of the cell in the channel resulted in the same deformation of the cell (data not shown).

The stress and strain states from the finite element simulations (Extended Data Fig. 4c) showed a pattern in stress magnitude consistent with analytical results (see above section). The axial strain increased monotonically from the upstream end to the downstream end along the cell length. The parameter ranges that matched experimental findings suggested greater turgor pressure in the bacteria submitted to larger extrusion loading (Supplementary Table Supplementary Table 3), which is in agreement with the conclusion from the analytical model (see above section).

The stress and strain states of bacteria under extrusion loading were compared to bacteria under agarose gel encapsulation (see "1.14. Agarose embedding assay of cell growth," Supplementary Table Supplementary Table 4). During elongation of a single cell, material is added to the cell envelope, thereby increasing cell length. When encapsulated in a stiff gel, the axial extension of the cell during growth is constrained (Extended Data Fig. 9a). The cell envelope is typically under tension due to turgor. The presence of the gel results in a reduction in axial tensile stresses in the bacterial cell envelope ($l_0$). As there is typically no expansion of *E. coli* in the hoop direction, hoop stress in an encapsulated cell remains the same as hoop stress in a free-floating cell. We modeled bacteria using a transversely isotropic linear elastic constitutive model (see section above) with applied turgor pressure and axial compressive forces $P_{gel}$ (applied on the cell poles as the cell grows). The resulting strain tensor is given in Eq. S22, where $\upsilon$ is the Poisson's ratio in the isotropic plane, $\upsilon_{ah}$ and $\upsilon_{ha}$ are the transverse direction Poisson's ratio of the cell envelope, $\sigma_{a,0}$ is the axial stress, $\sigma_{h,0}$ is the hoop stress, and $\sigma_{r,0}$ is the radial stress of a free-floating cell calculated from a thin-walled pressure vessel model.

$$\begin{pmatrix} \varepsilon_a \\ \varepsilon_h \\ \varepsilon_r \end{pmatrix} = \begin{pmatrix} 1/E_a & -\nu_{ha}/E_h & -\nu_{ha}/E_h \\ -\nu_{ah}/E_a & 1/E_h & -\nu/E_h \\ -\nu_{ah}/E_a & -\nu/E_h & 1/E_h \end{pmatrix} \begin{pmatrix} \sigma_{a,0} - P_{gel} \\ \sigma_{h,0} \\ \sigma_{r,0} \end{pmatrix} \qquad \text{Eq. S22}$$

Using a set of material properties that were consistent with extrusion loading experimental results (Supplementary Table Supplementary Table 3), we compared the cell envelope stress states during extrusion loading with those during gel encapsulation. The normal stresses generated by extrusion loading and gel encapsulation differ considerably (Supplementary Table Supplementary Table 4). Extrusion loading leads to increased axial stresses and reduced hoop stresses while gel encapsulation results in reductions in axial stress and negligible changes in hoop stress. Although the normal stresses are very different, decomposing the three-dimensional stress state into hydrostatic and octahedral shear components shows that both loading modes result in increases in octahedral shear stress (Fig. 3c-d, Extended Data Fig. 10).

**Supplementary Table 1.** Parameters for the cell envelope in the finite element model

| Parameter | Definition | Parameter Range | Reported Range |
|---|---|---|---|
| $E_a$ | Axial Young's modulus | 20 – 60 MPa | 20 – 150 MPa[6,7,11-15] |
| $P_{t,0}$ | Turgor pressure prior to extrusion loading | 100, 150, 200 kPa | 30 – 300 kPa[7,16,17] |
| $P_{t,load}$ | Turgor pressure during extrusion loading | $P_{t,0}$ – 500 kPa | – |
| $l_0$ | Unstressed cell length | 1000 – 2400 nm | – |

**Supplementary Table 2.** Applied pressure difference and cell dimensions for cells shown in Fig. 2a (with different $\Delta P$ binning)[a]

| $P_{ave}$ (kPa) | $\Delta P$ (kPa) | $n$ | $P_{u,ave}$ (kPa) | $P_{d,ave}$ (kPa) | $W_{ave} \pm SD$ (nm) | $L_{ave} \pm SD$ (nm) |
|---|---|---|---|---|---|---|
| 30.0 | 10 – 20 | 95 | 37.73 | 22.25 | 684 ± 165 | 2762 ± 438 |
| | 20 – 30 | 146 | 42.16 | 17.82 | 697 ± 134 | 2617 ± 467 |
| | 30+ | 296 | 47.28 | 12.70 | 645 ± 104 | 2596 ± 457 |

[a]Note: the parametric analysis identifies sets of unknown parameters (turgor pressure, Young's modulus, and cell length at unstressed state) that result in a cell width similar to what is seen experimentally. $P_{u,ave}$ is the average upstream pressure and $P_{d,ave}$ is the average downstream pressure for trapped cells. $W$ is the width of the cell expressed at mid-cell (halfway between the upstream and downstream ends of the cell) and $L$ is the length of the cell trunk (not including caps).

**Supplementary Table 3.** An example of parameter ranges using fixed and assumed values for the Young's modulus ($E_a$) and initial turgor pressure ($P_{t,0}$). Values were assumed from literature (see Reported Range in Supplementary Table Supplementary **Table 1**)

| $P_{ave}$ (kPa) | $\Delta P$ (kPa) | $E_a$ (MPa) | $P_{t,0}$ (kPa) | $P_{t,load}$ (kPa) | $l_0$ (nm) |
|---|---|---|---|---|---|
| 30.0 | 10 − 20 | 25 | 150 | 150 − 230 | 1400 − 2400 |
| | 20 – 30 | 25 | 150 | 190 − 280 | 1200 − 2000 |
| | 30+ | 25 | 150 | 260 − 300 | 1200 – 1600 |

**Supplementary Table 4.** The responses of stress and strain to increases in external loading of the two loading modes[a]

| **Loading mode** | $\sigma_a$ | $\sigma_h$ | $\sigma_r$ | $\tau$ | $\sigma_{hyd}$ | $\tau_{oct}$ | $\varepsilon_{hyd}$ | $\gamma_{oct}$ |
|---|---|---|---|---|---|---|---|---|
| Extrusion loading | ↑ | ↓ | ↓ | ↑ | − | ↑ | − | ↑ |
| Agarose gel encapsulation | ↓ | − | − | − | ↓ | ↑ | ↓ | − |

[a]Note: "−" denotes no change. The ↑ symbol indicates an increase in the specified stress or strain with an increase in the loading (either from greater $\Delta P$ or increased agarose content), ↓ is a decrease in stress or strain with increased loading. Stresses examined were axial ($\sigma_a$), hoop ($\sigma_h$), radial ($\sigma_r$), shear ($\tau$), hydrostatic ($\sigma_{hyd}$), and octahedral shear ($\tau_{oct}$). Strains examined were hydrostatic ($\varepsilon_{hyd}$) and octahedral shear ($\gamma_{oct}$).

#### 1.7. Bacterial strain construction

All strains used in this study were derived from the *E. coli* BW25113 strain (CGSC# 7739 Keio Collection, Yale; genotype: F-, Δ*(araD-araB)567*, Δ*lacZ4787*(::rrnB-3), $\lambda^-$, *rph-1*, Δ*(rhaD-rhaB)568*, *hsdR514*). The *cusA*$^{mE}$ (i.e., CALMF) strain was constructed and characterized in our previous work[4]. Briefly, CALMF (*cusA*-linker-mEos3.2-FLAG) was created via lambda Red recombineering, where a short, flexible linker L of 10 amino acids (sequence = AGSAAGSGEF) was used to connect mEos3.2-FLAG (a monomeric, irreversibly photoconvertible fluorescent protein mEos3.2[18,19] with a C-terminal FLAG tag) to the C-terminus of CusA at its chromosomal locus[4]. This CusA$^{mE}$ fusion protein is functional and stays intact, as shown by cell growth assays and Western blot[4].

#### 1.8. Cell sample preparation

**For single-molecule imaging:** CusA$^{mE}$ (i.e., CALMF) cells were grown in LB with chloramphenicol (25 µg/mL, USBiological) for 18 h in 37 °C with shaking (250 rpm). From this culture, a 1:100 dilution was prepared in LB containing chloramphenicol (25 µg/mL). This diluted 10 mL culture was incubated at 37 °C for 4 h (reaching $OD_{600}$ = 0.4) with shaking (250 rpm). The cells were centrifuged (4500 rpm, 4 °C) for 10 min. The supernatant was removed, and the resulting cell pellet was re-suspended in 10 mL of M9 medium supplemented with 8% v/v 50X MEM amino acids (GIBCO, cat. #: 11130051), 4% 100X MEM vitamins (GIBCO, cat. #: 11120052), and 0.4% glucose (Sigma-Aldrich, cat. #: G7528). The liquid suspension of cells was loaded into the microfluidic device as described earlier.

**For cell growth under copper stress in device:** For cell samples requiring copper stress (e.g., in measuring the rate of cell elongation and division), CALMF cell cultures were prepared as described above. The pellet was re-suspended in 10 mL of LB, and 25 µL of copper sulfate

(Mallinckrodt) was added to yield a 2.5 mM copper solution. This concentration of copper impacts *E.coli* cell growth but still renders the cells viable.

#### 1.9. Single-molecule tracking (SMT) and single-cell quantification of protein concentration (SCQPC)

SMT[20-26] via stroboscopic imaging and SCQPC were performed as previously described on an inverted fluorescence microscope (Olympus IX71; Supplementary Fig. 2)[2,4]. For SMT, the cells were first illuminated with a 405 nm laser (1–10 W/cm$^2$) for 20 ms to photoconvert a single mEos3.2 molecule (or none) from its green fluorescent form to its red fluorescent form. A series of 30 pulses of a 561 nm laser (21.7 kW/cm$^2$) in epi-illumination mode with pulse duration $t_{\mathrm{int}}$ = 4 ms and time lag $T_{\mathrm{tl}}$ = 60 ms were then used to excite the red mEos3.2. The resulting red mEos3.2 fluorescence was imaged by an EMCCD camera (Andor Technology, DU-897E-CSO-#BV), which was synchronized with the 561 nm laser pulses. This imaging scheme was repeated for 500 cycles for each field of view.

After the SMT step, SCQPC was performed on the same cells, in which the cells were illuminated with the 405 nm laser (1–10 W/cm$^2$) for 1 min to photoconvert all remaining mEos3.2 molecules, followed by 561 nm laser illumination for 2000 frames with the same laser power density and exposure time as in the SMT step to quantify the number of remaining mEos3.2 molecules. This 405-illumination and 561-excitation sequence was repeated once more to ensure all mEos3.2 molecules had photobleached. All CusA concentrations cited in the study correspond to those of CusA trimers, i.e., one-third of the total mEos3.2 concentration.

#### 1.10. Single-molecule imaging data analysis

The boundary of each cell inside tapered channels was determined from the optical transmission image. Due to the interference of tapered channels, automated detection of the cell boundary in the image was problematic, and a manual approach was used. An ellipse was hand-drawn to initially outline crudely each cell's boundary, which was then fitted with a cell-shape model of a cylinder with two hemispherical caps[2,27]. The resulting fitted cell boundary was then used as a mask to define the region of interest (ROI) for each cell in the corresponding fluorescence image. Fluorescence spots of individual CusA$^{\mathrm{mE}}$ molecules from SMT were identified within each ROI [i.e., if the pixel intensities of these spots were greater than a user-defined threshold (the mean pixel intensity value + 4 standard deviations of the whole image)], fitted with a 2-D Gaussian point spread function, and position-localized to nanometer precision (typically ~40 nm precision), as previously described[2].

To obtain more precise measurements of the boundary of each cell, we took horizontal and vertical line profiles of each cell's optical transmission image (Supplementary Fig. 3). For the horizontal line profile, the midpoints of each of the two curves surrounding the cell were used to determine the cell length. For the vertical line profile, the midpoints of the two taper channel walls were used to determine the width. In cases of ambiguity, the lengths and widths obtained from line profiling were each dilated by one pixel (135.4 nm) in each direction. From these refined measurements, the lengths and widths of the cells can be accurately reported (Supplementary Table 5). The more precise measurements of cell lengths and widths were used to generate refined cell boundaries (i.e., new ROIs), which were used to clean up the fluorescence localizations of individual $CusA^{mE}$ molecules. Fluorescence localizations outside these new ROIs were eliminated as false hits.

We further examined the cells with higher numbers of detected trajectories (i.e., the top 5% cells). If the cell dominantly had fluorescence localizations that appear to stay at the same location (i.e., within ~270 nm, which is about diffraction-limited spot size) repetitively for long periods (i.e., for an average of 8 minutes), the cell was removed, as these apparent stationary bright objects are likely from dust particles. For all 1337 cells analyzed, a total of 10 cells (i.e., less than 1%) were discarded due to the presence of these dust-particle-like behaviors.

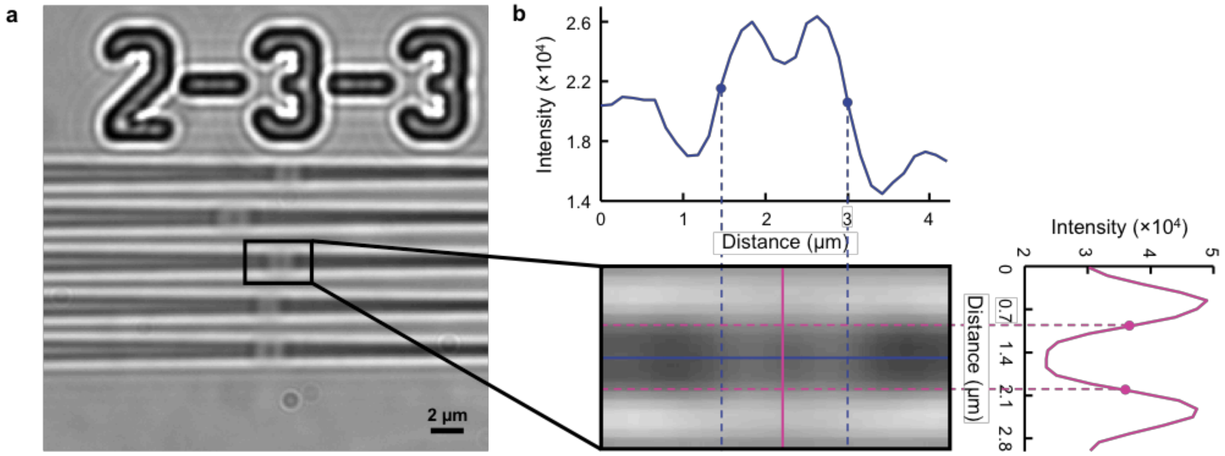

**Supplementary Figure 3. Line profiling for determining accurate cell boundary. a**, Exemplary transmission image of cells trapped in tapered channels. **b**, Horizontal (blue) and vertical (pink) line profiling of a cell from a (lower left). For the horizontal line profile (upper), the midpoints (blue dots) of each of the two curves surrounding the cell were used to determine the cell length. For the vertical line profile (right), the midpoints (pink dots) of the two taper channel walls were used to determine the cell width.

**Supplementary Table 5.** Applied pressure difference and cell dimensions for cells shown in Fig. 2c-f[a]

| $P_{ave}$ (kPa) | $\Delta P$ (kPa) | $n$ | $L_{ave} \pm SD$ (μm) | $W_{ave} \pm SD$ (μm) |
|---|---|---|---|---|
| All | 5.6 ± 3.3 | 481 | 2.14 ± 0.49 | 0.91 ± 0.19 |
| | 15.1 ± 2.4 | 438 | 2.24 ± 0.49 | 0.67 ± 0.12 |
| | 32.1 ± 4.8 | 408 | 2.29 ± 0.46 | 0.62 ± 0.08 |
| 12.5 | 4.9 ± 2.2 | 205 | 2.04 ± 0.51 | 1.01 ± 0.16 |
| | 11.5 ± 1.4 | 199 | 2.22 ± 0.44 | 0.71 ± 0.13 |
| | 15.5 ± 0.7 | 188 | 2.22 ± 0.53 | 0.63 ± 0.08 |
| 30.0 | 4.3 ± 3.4 | 189 | 2.24 ± 0.47 | 0.88 ± 0.16 |
| | 18.8 ± 4.1 | 194 | 2.27 ± 0.48 | 0.72 ± 0.13 |
| | 33.4 ± 3.6 | 349 | 2.28 ± 0.45 | 0.61 ± 0.07 |

[a]: $P_{ave}$ denotes the average pressure applied to the cells; $\Delta P$ is the pressure difference experienced across the cell; $n$ represents the number of cells; $L$ is the length of the cell as calculated from line profiling; $W$ is the width of the cell expressed at mid-cell (halfway between the upstream and downstream ends of the cell) as calculated from line profiling; and SD is standard deviation.

#### 1.11. Quantification of stage drift: it is insignificant within our experimental time.

To check and ensure that our microscope stage did not drift significantly during our imaging — each fluorescence imaging movie was acquired over ~20 minutes, consisting of SMT (~15 min) and 2 rounds of SCQPC (~2 min each), we took the optical transmission images of our device containing the cells before and after each movie was recorded (Supplementary Fig. 4a). Our device had numerical labels etched in, and each transmission image contained at least one fully-intact symbol from these labels (number or hyphen), which acted as reference position markers. Upon selecting the transmission image of interest, the Canny edge detection algorithm in MATLAB was used to determine all of the edges in the entire image (Supplementary Fig. 4b). A mask was applied to include all the edges surrounding the position marker of interest. All connected regions within this mask were filled, and the region with the maximum number of pixels (i.e., greatest area) was identified (Supplementary Fig. 4c). The centroid position (geometric center) of this identified region was calculated (Supplementary Fig. 4c-d). If multiple symbols were present in one transmission image, multiple centroids were determined, and their averages were used as position markers.

Upon analyzing the hundreds of centroid positions over many imaging areas, the average stage drifts in the $x$ and $y$ directions are 29.3 ± 27.6 nm and 44.3 ± 37.8 nm, respectively, over the duration of one movie (~20 min), corresponding to average drift rates of 0.28 ± 0.028 and 0.43 ± 0.034 nm $s^{-1}$ (Supplementary Fig. 4e). Considering the localization precision of our single-molecule imaging is ~40 nm, the magnitude of stage drift falls within the localization error. Thus, no subsequent steps were needed to correct for stage drift in analyzing the single-molecule tracking trajectories.

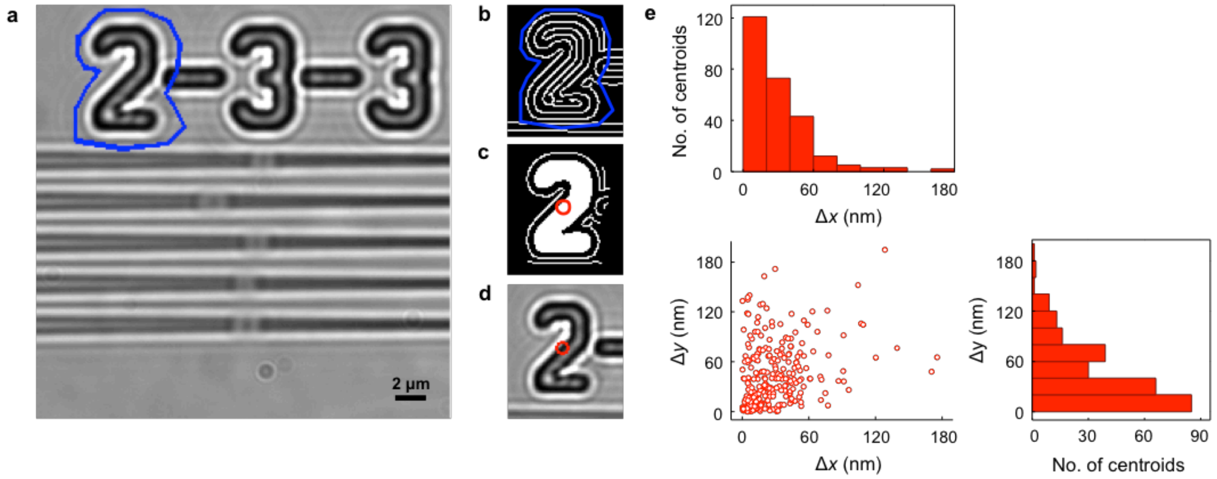

**Supplementary Figure 4. Edge detection and centroid determination.** **a**, Transmission image containing the taper set label "2−3−3" etched into the imaging wafer. Blue dots connected by white lines indicate the selection of the mask for the desired symbol (position marker). **b**, The Canny edge detection algorithm selects the edges of the entire transmission image. The intensity values are converted into binary. **c**, Selection of the edges within the mask indicated in panel b, followed by filling in the connected regions. The filled region with the maximum number of pixels (i.e., the greatest area) — i.e., the solid white innermost region — is identified. The centroid position (i.e., the geometric center) of this identified region is calculated, depicted here by the red superimposed circle. This process was repeated for each fully-intact symbol on each set of transmission images taken before and after the recording of each movie. **d**, Overlay of the centroid position (red circle) determined from c on the original transmission image of a. **e**, Histograms (upper left and bottom right) and scatter plot (lower left) of the stage drift in $x$ and $y$ direction for all analyzed centroids.

### 1.12. Analysis of diffusive behaviors of CusA$^{mE}$ in cells to determine its diffusive states and the associated diffusion constants and fractional populations

From SMT data, the diffusive behaviors of CusA$^{mE}$ can be extracted by analyzing the probability distribution function (PDF) of the displacement length $r$ per time lapse ($T_{tl}$) of each tracked CusA$^{mE}$ molecule, as we reported in our previous study of CusCBA assembly[4]. Only the first displacement was used here from each tracking trajectory to avoid biasing toward long trajectories[2]. The experimentally determined histogram of $r$ (i.e., un-normalized PDF of $r$) was initially transformed via an inverse transformation method ITCDD[4,28,29] to deconvolute out the cell confinement effect that distorts the displacement distribution and was then fitted by a linear combination of two terms, each representing a diffusive state that follows the Brownian diffusion model (Eq. S23) (the actual fitting was done on the integrated form of the PDF to be numerically more robust):

$$\mathrm{PDF}(r,T_{\mathrm{tl}})_{\mathrm{unnormalized}} = N\left[\frac{A_{\mathrm{m}}r}{2D_{\mathrm{m}}T_{\mathrm{tl}}}\exp\left(-\frac{r^2}{4D_{\mathrm{m}}T_{\mathrm{tl}}}\right)+\frac{A_{\mathrm{s}}r}{2D_{\mathrm{s}}T_{\mathrm{tl}}}\exp\left(-\frac{r^2}{4D_{\mathrm{s}}T_{\mathrm{tl}}}\right)\right] \qquad \text{Eq. S23}$$

where $N$ is a scaling factor, $D$ is diffusion constant, $A$ is fractional population, and the subscripts m and s refer to the mobile (faster) and stationary (slower) populations, where $A_{\mathrm{m}} + A_{\mathrm{s}} = 1$. This analysis was applied to all PDF($r$)'s across all experimental conditions to evaluate how the fractional populations and diffusion constants of the two states varied.

Upon ITCDD transformation of the original experimental PDF, the resulting distribution of displacement lengths are fitted by the Brownian diffusion model as in Eq. S23 above. However, we often observe that after ITCDD, there are some displacement lengths that are too large to be reasonable for a membrane protein[4] and need to be discarded. To determine the appropriate upper limit of displacements for the transformed PDF, we performed simulations of a statistically-saturated number of diffusion trajectories (5000) with a given diffusion constant on the membrane of a cell matching the average cell geometry of our experimental data. The resulting simulated displacements were transformed via ITCDD, from which the diffusion constant was extracted from Brownian diffusion model analysis. Each simulation was repeated four times for each diffusion constant input (diffusion constant inputs of 0.1 to 1.1 μm$^2$ s$^{-1}$ were sampled). The extracted diffusion constants from the ITCDD-transformed simulations very well recover the intrinsic diffusion constant input, validating our ITCDD method (Supplementary Fig. 5a), which we showed previously in our method paper[28]. Next, the displacement distributions from the simulations were truncated in 0.1 μm increments at the 95% population of the theoretical PDF (i.e., the displacement threshold $r_{\mathrm{upper}}$ @ 95%). We chose 95% confidence to be better than our typical experimental uncertainty. This established our calibration curve for the upper-limit truncation on the ITCDD-transformed PDF (Supplementary Fig. 5b, red points). We then iteratively truncated each of our ITCDD-transformed experimental PDF (going from high to low upper limit) until the value of the extracted experimental diffusion constant of the mobile faster state agreed with the expected diffusion constant from the calibration curve at the same truncation $r_{\mathrm{upper}}$ (i.e., crossed the red calibration curve, Supplementary Fig. 5b). The PDF analysis results at this $r_{\mathrm{upper}}$ value were subsequently taken as our final results.

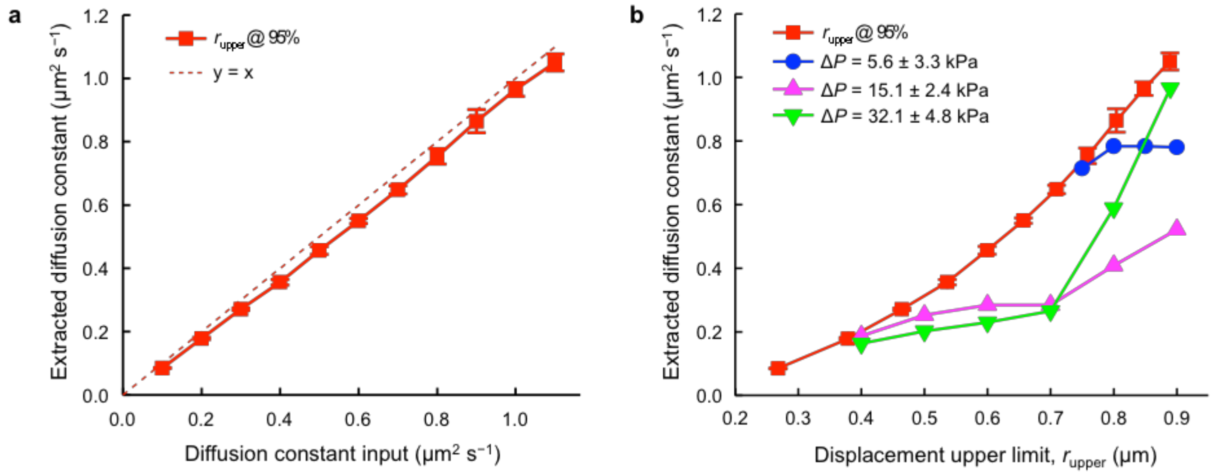

**Supplementary Figure 5. Truncation of ITCDD-transformed PDF.** **a**, Diffusion constant input plotted against extracted diffusion constant fitted from simulated displacements that were transformed via ITCDD and truncated at 95% population inclusion. Each red square represents the average extracted diffusion constant from four simulations; y-error bars are s.d. For reference, a diagonal line ($y = x$) was plotted to illustrate that the values of the extracted diffusion constants from the simulations very closely align with the original diffusion constant inputs. **b**, The $r_{upper}$ value at 95% of ITCDD-transformed PDF vs. the extracted diffusion constant (red points). The blue, pink, and green curves represent the three $\Delta P$ groups of the experimental data. The ITCDD-transformed PDF from each group was truncated iteratively in the direction of high to low $r_{upper}$ (indicated by the direction of the color-corresponding arrows) until the extracted diffusion constant meets the red calibration curve. The corresponding upper limit of displacement (as indicated by each color-corresponding vertical dashed line) was used for subsequent analysis of the ITCDD-transformed PDF.

### 1.13. Determination of effective disassembly rate constant

The disassembly rate constant of CusA from the CusCBA complex was determined by analyzing the distribution of microscopic residence times $\tau$ of single CusA$^{mE}$ molecules at the stationary state (i.e., assembled state) (Extended Data Fig. 6a-e); these single-molecule residence times were obtained by up-thresholding the displacement vs. time trajectories. Two different methods are described below.

In the first method (Method 1), the displacement threshold $r_0$ (≈100 nm) was determined from the resolved distribution of displacements (Extended Data Fig. 6d), as described in our previous work[4]. Specifically, $r_0$ was determined by determining the intersection between PDF($r$) of the two slower states before inverse transformation (these two slower states correspond to the stationary and mobile states after inverse transformation while the third (fastest) state with apparent diffusion constant of 4.70 ± 0.91 μm$^2$ s$^{-1}$ is an artifact from the cell confinement effect as we showed previously[4,28]: $r_0$ = 99 nm for $\Delta P$ = 5.6 ± 3.3 kPa; $r_0$ = 98 nm for $\Delta P$ = 15.1 ± 2.4 kPa; $r_0$ = 95 nm for $\Delta P$ = 32.1 ± 4.8 kPa.

In the second method (Method 2), $r_0$ (= 127 nm) was selected as to include 99% of the population of the stationary state ($D$ = 0.015 ± 0.001 μm$^2$ s$^{-1}$ for $\Delta P$ = 1.1 to 38.0 kPa) (Extended Data Fig. 6d). Note the displacement distribution here is before applying the inverse transformation method, but the diffusion constant of the stationary state is essentially the same as that after the inverse transformation ($D_s$ = 0.03 to 0.04 μm$^2$ s$^{-1}$; Fig. 2d), as the cell confinement effect only significantly distorts the displacement distribution of the faster mobile state.

For both methods of determining $r_0$, we obtained the thresholded individual residence times (Extended Data Fig. 6b-c), and their distributions $f(\tau)$ (e.g., Extended Data Fig. 6e) were fitted with the following equation (Eq. S24):

$$f(\tau) = N \exp\left[-\left(k_{\mathrm{bl}}\frac{T_{\mathrm{int}}}{T_{\mathrm{tl}}} + k_{\mathrm{d}}\right)\tau\right] \qquad \text{Eq. S24}$$

where $N$ is a scaling parameter, $k_{\mathrm{bl}}$ is the photobleaching/blinking rate constant of mEos3.2 (independently determined by analyzing the distribution of the mEos3.2 fluorescence on-times in the SMT trajectories, as described in our previous work[2]; $k_{\mathrm{bl}}$ = 256 ± 14 s$^{-1}$; 273 ± 16 s$^{-1}$; 268 ± 12 s$^{-1}$ for the three groups in Fig. 2f and Extended Data Fig. 6f), $T_{\mathrm{int}}$ is the laser integration time (4 ms), $T_{\mathrm{tl}}$ is the time lapse (60 ms), and $k_{\mathrm{d}}$ is the effective first-order disassembly rate constant[4].

#### 1.14. Agarose embedding assay of cell growth

Our experimental procedure for the agarose embedding assay to probe mechanical effects on cell growth was adapted from Auer et al.[12].

CusA$^{\mathrm{mE}}$ (i.e., CALMF) cells were grown in LB with chloramphenicol (25 μg/mL, USBiological) for 18 h in 37 °C with shaking (250 rpm). From this culture, a 1:100 dilution was prepared in LB with chloramphenicol (25 μg/mL). This diluted 10 mL culture was incubated at 37 °C for 4 h (reaching $OD_{600}$ = 0.4) with shaking (250 rpm). The cells were centrifuged (4500 rpm, 4 °C) for 10 min. The supernatant was removed, and the resulting cell pellet was re-suspended in 1 mL LB.

UltraPure™ agarose (Invitrogen) was prepared as solutions of 0% (control), 0.25%, and 0.5% w/v in 20 mL LB, corresponding to 0, 2.5, 5 mg/mL, respectively. *Note, it is not possible to prepare agarose gel at concentrations in lower than <0.25%, as in this concentration agarose gel does not solidify.* The agarose solutions were heated in a microwave until the agarose was fully dissolved. The solutions were then transferred to a 50 °C water bath, where they were incubated for at least 20 min to ensure that the temperature of the agarose solution was equilibrated, and to keep the agarose solution in liquid state for the following steps. To prepare each sample for spectrophotometric analysis, 2 mL of this agarose solution was pipetted into a cuvette, and 2 μL of chloramphenicol (from a 25 mg/mL stock, USBiological) was added. The appropriate amount of $CuSO_4$ (Mallinckrodt) was subsequently added to make final copper concentrations of 0, 0.3, 0.6, and 1 mM. Finally, 67 μL of cells were added to each cuvette. The contents of the cuvette were then gently mixed (resuspended) thoroughly via pipetting; each cuvette was then sealed with Parafilm (Bemis NA), and the cuvettes were allowed to cool to room temperature, where the agarose solution could solidify (typically takes ~45 min).

Once all samples were prepared, the cuvettes were shaken (250 rpm) at room temperature for a period of 5 hr (a time by which all samples had experienced their maximum rate of growth). The absorbance (λ = 600 nm) for each sample was measured (Beckman Coulter DU800 UV-Vis spectrophotometer) against its corresponding blank every 30 minutes over this 5 hr duration. The maximum rate of growth was determined for each condition (Extended Data Fig. 9b), which was used by Auer et al.[12] to report mechanical effects on cell growth rate. The *sensitivity* of cells to copper upon agarose stress was also determined by computing the change in maximum growth rate relative to 0 mM Cu (Extended Data Fig. 9c-d).

### 2. Pairwise distance distribution analysis supports that clustering or declustering is not responsible for the observed population shifts in CusA disassembly

In our previous work, we discovered CusA$^{mE}$ trimers tend to cluster together under certain conditions (e.g., when examined in a Δ*cus*CB double-deletion strain)[4]. To determine whether CusA clustering played a role in the increased disassembly observed when cells are exposed to increased levels of mechanical stress (i.e., Δ*P*), we examine the pairwise distance distributions of individual CusA$^{mE}$ proteins that were tracked at different pressure conditions. Pairwise distance distribution (PWD) is defined here as the range of Euclidean distances between pairs of first localizations of individual CusA$^{mE}$ proteins. The more prominent the clustering, the shorter the pairwise distances are expected.

Extended Data Figs. 5a-b show the normalized pairwise distance distributions for the three Δ*P* groups of individual cells (>400 cells per pressure condition/group), and their differences, respectively. Within experimental uncertainty, the distributions are indistinguishable from one another, and no significant shortening in the pairwise distances was observed with increasing Δ*P*, indicating that clustering of CusA trimers does not play a significant role in the observed increase in the CusA disassembly upon greater mechanical stress. Note in Extended Data Fig. 5b, we consider the small peak at ~70 nm in the pink curve to be insignificant, as it is close to the uncertainty (i.e., standard deviation ~57 nm) of the pairwise distances.

### 3. Stiffness–sorted analysis suggests softer cells are more responsive to pressure–induced deformation)

To evaluate whether cell stiffness is related to mechanically inducible CusCBA disassembly, we used the distance traveled by the cell in the tapers as an approximate metric for cell stiffness[1] — the less stiff the cell, the farther the cell can travel into the taper. Upon plotting each cell's distance traveled against its corresponding pressure difference Δ*P*, we fitted the data with a saturation curve (Supplementary Fig. 6a).

We then computed each cell's residual by taking the difference between the cell's actual distance traveled and its predicted value from the saturation curve equation (Supplementary Fig. 6b). To account for uncertainty, we removed cells with residuals between −3 and +3 μm; the range was chosen to be significantly larger than the average length of the cells studied. The resulting ($n$ = 1011) cells were sorted into two categories: one category of cells with positive residuals and presumably "less stiff," and the other with negative residuals and "more stiff."

For each category, we further sorted the cells into three groups of similar Δ*P* values, and analyzed the CusA$^{mE}$ diffusive behaviors in each group The resulting fractional population of the disassembled state suggests that less stiff cells show perhaps more response to pressure-induced CusCBA disassembly (Supplementary Fig. 6c).

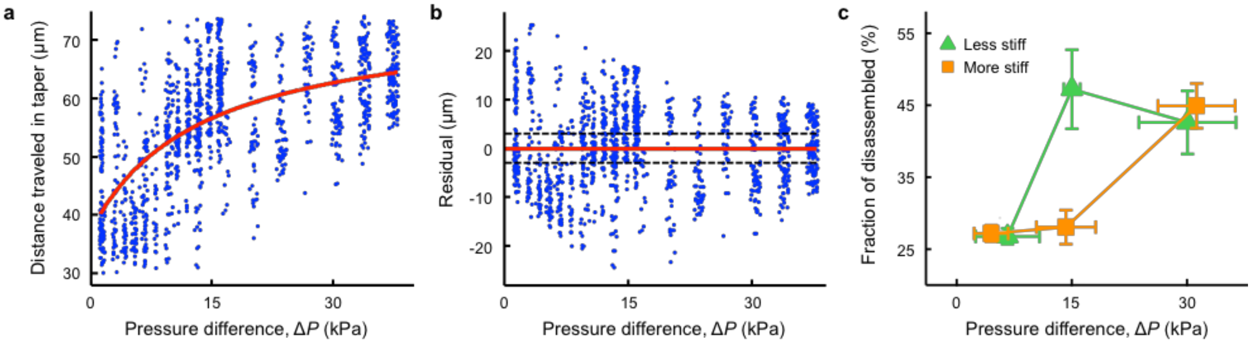

**Supplementary Figure 6. Stiffness-sorted analysis suggests less stiff cells are more responsive to pressure-induced CusCBA disassembly. a**, Scatter plot of distance traveled in the tapered channels as a function of pressure difference ($\Delta P$). Each blue dot represents one cell. Red line denotes saturation curve fitting: Distance traveled = $[74.31(\Delta P) + 511.2]/(13.75 + \Delta P)$. **b**, Scatter plot of each cell's residual vs. pressure difference. Red line represents the saturation curve; the black dashed lines are at residuals of +3 and −3 μm. All cells within these two bounds were eliminated from the analysis presented in panel **c**. **c**, Fractional population of the disassembled state of CusA with increasing $\Delta P$ for cells of differing stiffness. Green triangles denote cells with positive residuals (less stiff cells); orange squares denote cells with negative residuals (more stiff cells).
